# Major sex differences in allele frequencies for X chromosome variants in the 1000 Genomes Project data

**DOI:** 10.1101/2021.10.27.466015

**Authors:** Zhong Wang, Lei Sun, Andrew D. Paterson

## Abstract

An unexpectedly high proportion of SNPs on the X chromosome in the 1000 Genomes Project phase 3 data were identified with significant sex differences in minor allele frequencies (sdMAF). sdMAF persisted for many of these SNPs in the recently released high coverage whole genome sequence, and it was consistent between the five super-populations. Among the 245,825 common biallelic SNPs in phase 3 data presumed to be high quality, 2,039 have genome-wide significant sdMAF (p-value <5e-8). sdMAF varied by location: (NPR)=0.83%, pseudo-autosomal region (PAR1)=0.29%, PAR2=13.1%, and PAR3=0.85% of SNPs had sdMAF, and they were clustered at the NPR-PAR boundaries, among others. sdMAF at the NPR-PAR boundaries are biologically expected due to sex-linkage, but have generally been ignored in association studies. For comparison, similar analyses found only 6, 1 and 0 SNPs with significant sdMAF on chromosomes 1, 7 and 22, respectively. Future X chromosome analyses need to take sdMAF into account.

---

After the striking observation that the X chromosome was excluded from most genome-wide association studies (GWAS) (1), there has been a slow increase in the incorporation of the X chromosome. Several association methods have recently been developed for the X chromosome (2-6). However, these X chromosome specific downstream methods, similar to those developed for the autosomes, typically presume high quality data and implicitly assume that there is no sex difference in minor allele frequency (sdMAF).

Most genotype calling, imputation and sequence analyses of X chromosomal variants apply methods and tools that were developed for the autosomes (1, 3). However, there are reports that genotype missing rate is higher for SNPs on the X chromosome than autosomes (1, 3). In the non-pseudoautosomal region (NPR) of the X chromosome males are hemizygous, meaning that the intensity of allele signals from genotyping arrays, or the number of reads from sequencing, is half that of females. This may result in variant positions having higher missing rates for males than females.

In addition to higher missing rates for variants on the X chromosome, the two pseudo-autosomal regions (PARs), PAR1 and PAR2, create further challenges for the analysis at the boundaries between PARs and NPR. Because there is no recombination of NPR, but PAR1 and PAR2 recombine, variants in PARs close to the PAR-NPR boundaries are linked to variants in NPR of the X and Y chromosomes in a sex-specific fashion. Although the effects of sex-specific recombination rates in PAR1 and PAR2 on linkage have been examined for non-parametric linkage analysis of affected sibpairs (7), the implications for X chromosomal data collected for association studies have not been well explored. The controversial XTR/PAR3 in Xq21.3 (8) adds further complexity as it is embedded in the non-pseudoautosomal region.

Recently, sex-differences in allele frequency in PAR1 and PAR2 were described using the African super-population from the phase 3 data of the 1000 Genome Project (9), but the rest of the X chromosome and other four super-populations of the 1000 Genome Project were not examined. Evolutionary dynamics, including recombination within the PAR regions, were reasoned as a major contributing factor to sdMAF, but genotyping errors or the agreement with the high coverage data were not examined.

We hypothesize that sex difference in MAF exists across the X chromosome in human populations, including NPR and at the boundaries between NPR and PAR1, PAR2 and PAR3, and it is more prevalent in low-coverage whole genome sequence data. We use publicly available phase 3 data from the 1000 Genomes Project to test for sdMAF across the five super-populations. We compare the results with the recently released high coverage whole genome sequence to determine whether genotyping error is contributory. Better understanding of the different sources of sdMAF is critical to developing X chromosome-suitable analytical strategies, from improved data collection and imputation to more robust association methods for variants on the X chromosome.

## Results

The proportions of males and females were similar across all of the 26 populations of the 1000 Genomes Project (Figure S1 in the Supplementary Information).

### Phase 3 X chromosome-wide sdMAF

For biallelic SNPs with global MAF≥5% (Figure 1), the Manhattan plot for sdMAF p-values (Figure 2A) shows that a non-negligible proportion of X chromosome SNPs have significant sex difference in MAF even at the genome-wide significance threshold of p-value <5e-8: 0.83% of SNPs in NPR, 0.29% in PAR1, 13.1% in PAR2, and 0.85% in PAR3. The excesses of small p-values for sdMAF testing in all four regions of the X chromosome are also evident from both the QQ plots (Figure S2) and histograms of the sdMAF p-values (Figure S3). SNPs with significant sdMAF are located across the X chromosome but tend to cluster in specific regions (Figure 2A).

**Figure 1.**
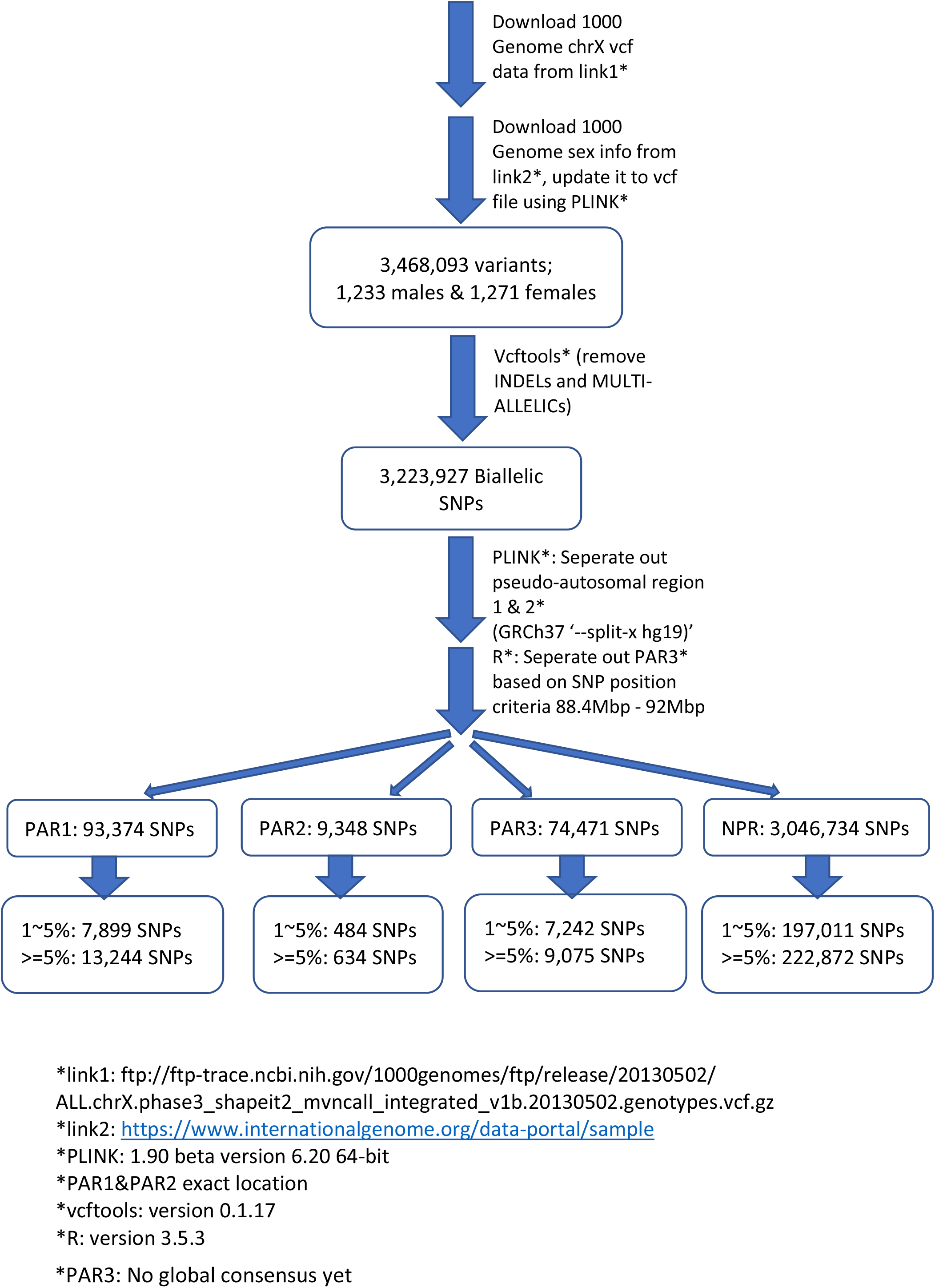
Pipeline for selection of X chromosome biallelic SNPs with global MAF ≥5%, presumed to be of high quality, from the 1000 Genomes Project phase 3 data on GRCh37. Variants were placed into the NPR, PAR1, PAR2, and PAR3 regions based on positions available from The Genome Reference Consortium and (8). For detailed counts of variant types and global MAF by regions, see Table S1 in the Supplementary Information.

**Figure 2.**
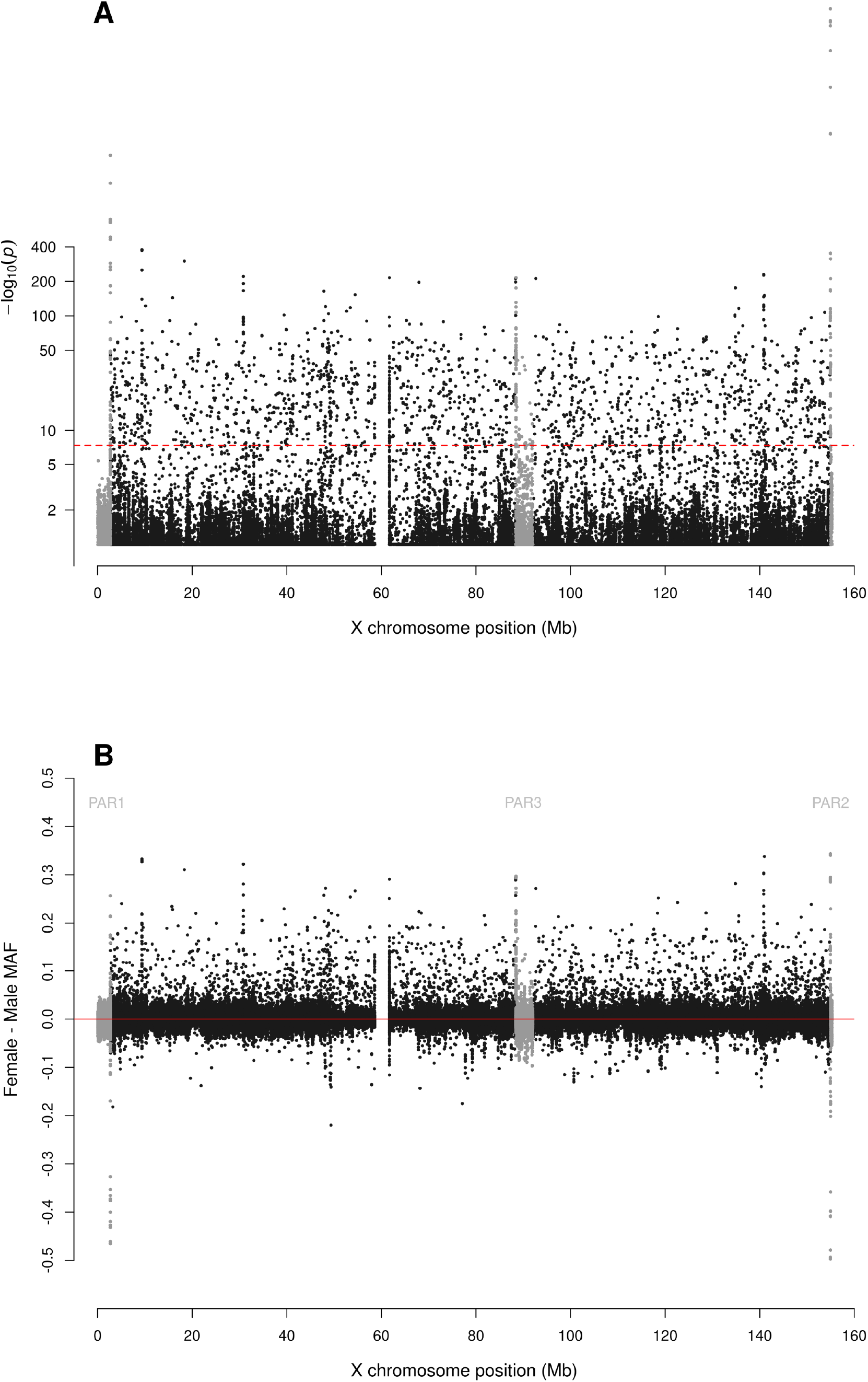
Manhattan plot for testing for sex difference in MAF across the X chromosome from the 1000 Genomes Project phase 3 data on GRCh37. A: sdMAF p-values for bi-allelic SNPs with global MAF ≥5% presumed to be of high quality. SNPs in the PAR1, PAR2 and PAR3 regions are plotted in grey, with PAR3 located around 90 Mb. Y-axis is −log10(sdMAF p-values) and p-values >0.1 are plotted as 0.1 (1 on −log10 scale) for better visualization. The dashed red line represents 5e-8 (7.3 on the −log10 scale). B: Female - Male sdMAF for the same SNPs in part A. For Zoomed-in plots for the PAR1, PAR2 and PAR3 regions see Figures 3, 4, and 5, respectively.

### Regional variation in sdMAF

Figure 2B shows that the direction of the sex difference in MAF varies by genomic location. Of the SNPs with significant sdMAF we observed females generally having higher MAF than males: NPR=93%, PAR2=59% and PAR3=86% among the SNPs with genome-wide significant sdMAF, with the exception of PAR1 (31%). Specifically, females have higher MAF at around 30 Mb (GRCh37) and at the q-arm of the centromere (Figure 2B), as well as at the centromeric boundary of PAR3 (Figure 5). In contrast, at the region of PAR1 close to the NPR boundary, males tend to have higher MAF among the SNPs with significant sdMAF (Figure 3). Finally, there are sets of PAR2 SNPs close to the NPR boundary that have higher MAF in females, while other sets of SNPs have higher MAF in males (Figure 4).

**Figure 3.**
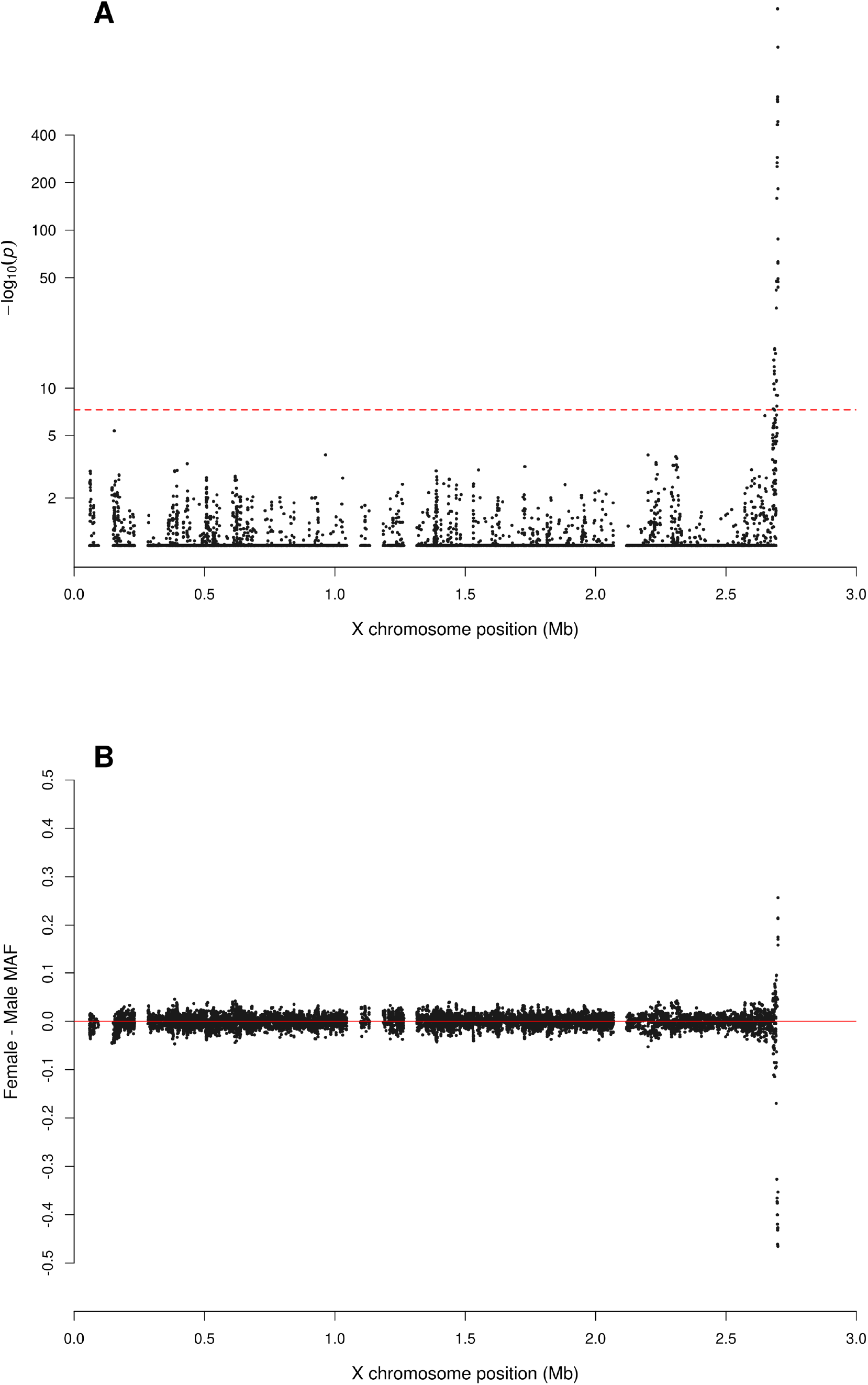
Zoomed-in plot for testing for sex difference in MAF across PAR1 of the X chromosome from the 1000 Genomes Project phase 3 data on GRCh37. A: sdMAF p-values for bi-allelic SNPs with global MAF ≥5% presumed to be of high quality. Y-axis is −log10(sdMAF p-values) and p-values >0.1 are plotted as 0.1 (1 on −log10 scale) for better visualization. The dashed red line represents 5e-8 (7.3 on the −log10 scale). B: Female – Male sdMAF for the same SNPs in part A, clearly showing PAR1 SNPs with significant sdMAF tend to cluster at the NPR-PAR1 boundary around 2.6 Mb.

**Figure 4.**
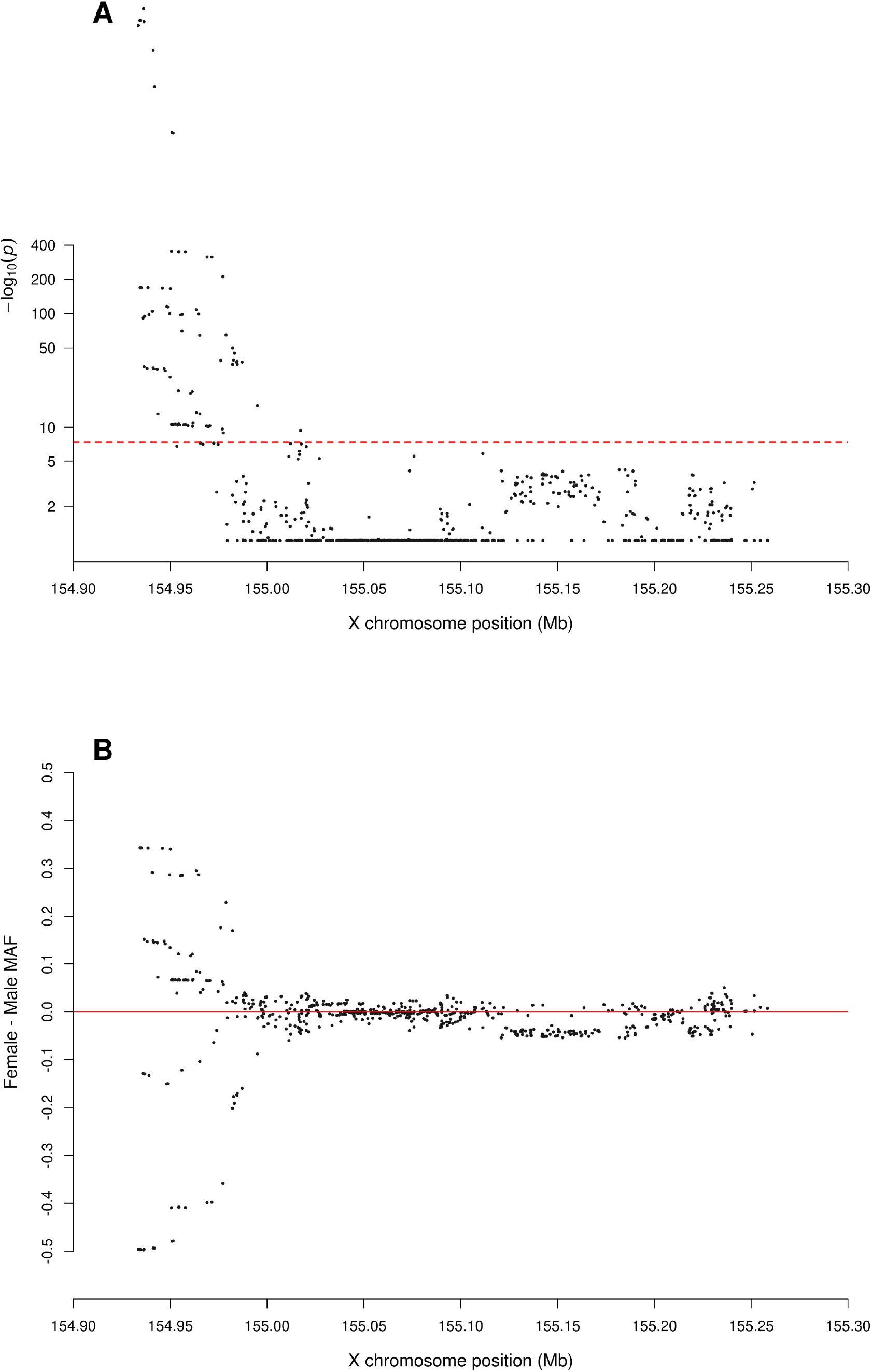
Zoomed-in plot for testing for sex difference in MAF across PAR2 of the X chromosome from the 1000 Genomes Project phase 3 data on GRCh37. A: sdMAF p-values for bi-allelic SNPs with global MAF ≥5% presumed to be of high quality. Y-axis is −log10(sdMAF p-values) and p-values >0.1 are plotted as 0.1 (1 on −log10 scale) for better visualization. The dashed red line represents 5e-8 (7.3 on the −log10 scale). B: Female – Male sdMAF for the same SNPs in part A, clearly showing PAR2 SNPs with significant sdMAF tend to cluster at the NPR-PAR2 boundary around 88.5 Mb.

**Figure 5.**
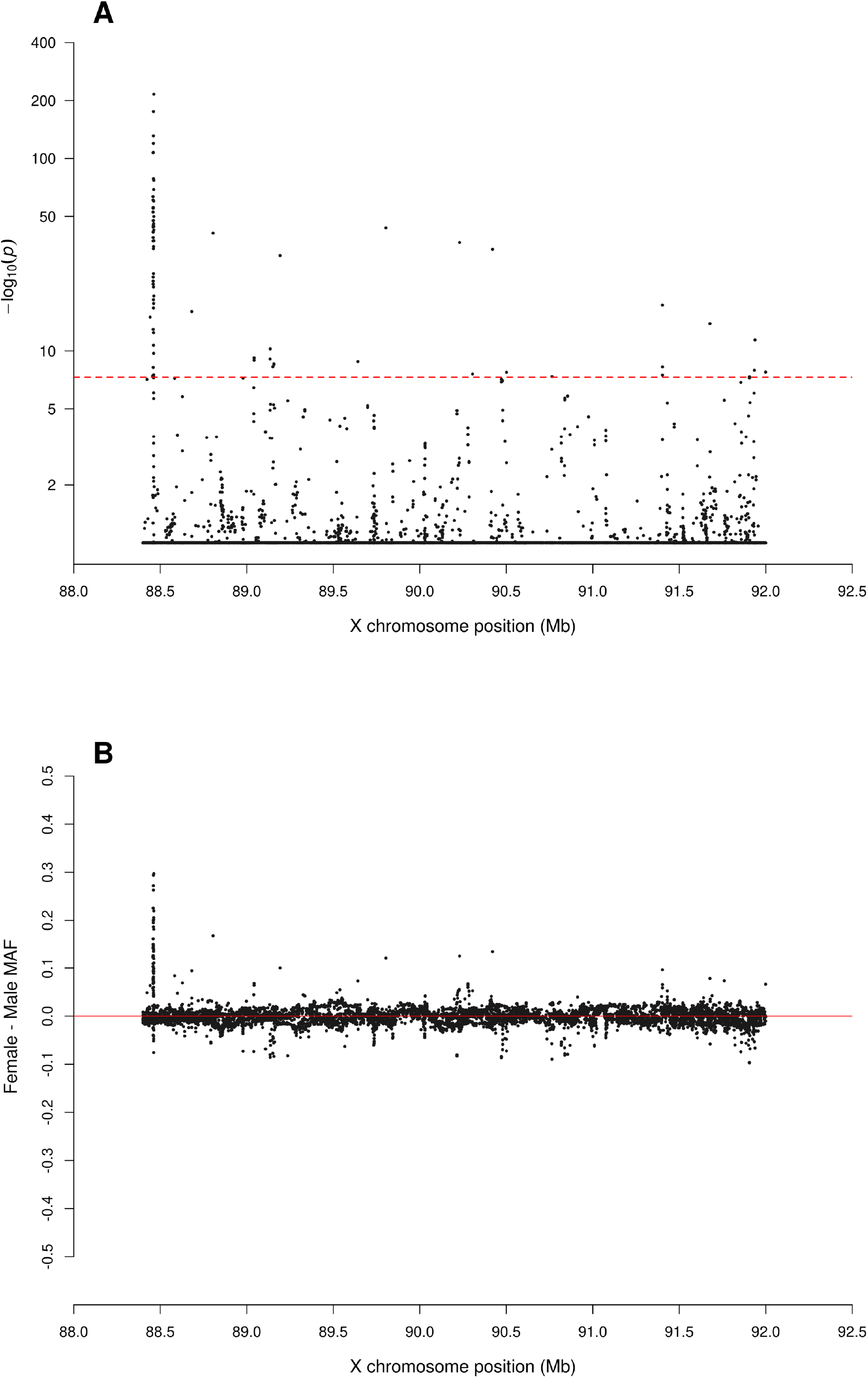
Zoomed-in plot for testing for sex difference in MAF across PAR3 of the X chromosome from the 1000 Genomes Project phase 3 data on GRCh37. A: sdMAF p-values for bi-allelic SNPs with global MAF ≥5% presumed to be of high quality. Y-axis is −log10(sdMAF p-values) and p-values >0.1 are plotted as 0.1 (1 on −log10 scale) for better visualization. The dashed red line represents 5e-8 (7.3 on the −log10 scale). B: Female – Male sdMAF for the same SNPs in part A, clearly showing PAR3 SNPs with significant sdMAF tend to cluster at one of the NPR-PAR3 boundaries around 88.5 Mb.

We then examined how the sdMAF relates to the sex-combined MAF. Figure 6 provides Bland-Altman plots (10) separately for each of the four regions. Of note, for NPR and PAR3 (Figures 6A and 6D), SNPs with significant sdMAF tend to have higher MAFs in females and predominantly had sex-combined MAFs in the range 25%-40%. In contrast, for PAR1 (Figure 6B), SNPs with significant sdMAF tend to have higher MAFs in males. Finally, for PAR2 (Figure 6C), there are sets of clustered SNPs with significant sdMAF in either direction (Figure 6C).

**Figure 6.**
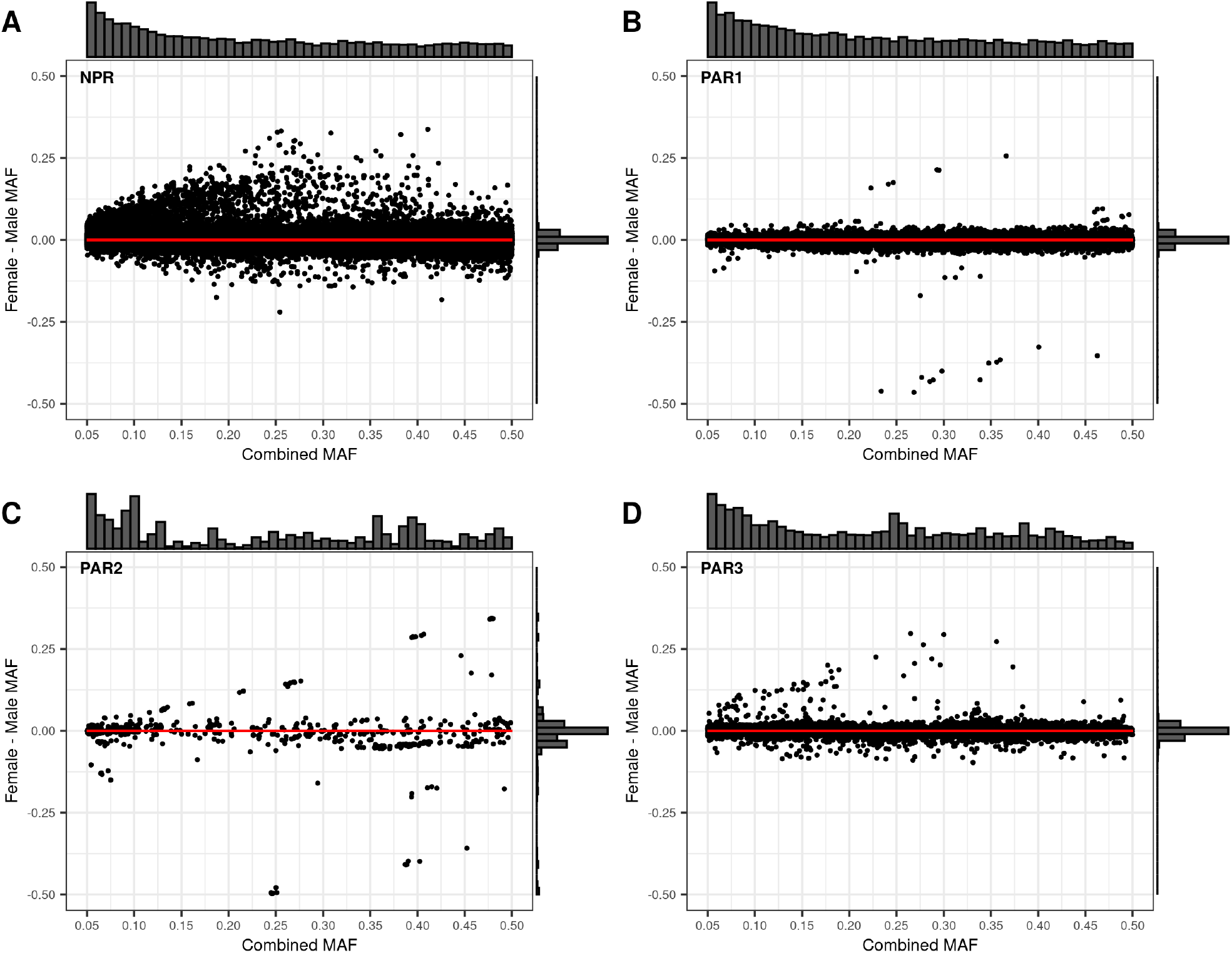
Bland-Altman plots comparing the Female - Male sex difference in MAF to the sex-combined MAF across the X chromosome from the 1000 Genomes Project phase 3 data on GRCh37. Regions are plotted separately A: NPR; B: PAR1, C: PAR2; D: PAR3. For each of the four regions, the histogram at the top of the Bland-Altman plot shows the distribution of the sex-combined MAF for bi-allelic SNPs with global MAF ≥5% presumed to be of high quality. The histogram to the right of the plot shows the distribution of the Female - Male sdMAF.

### In-depth analysis of eight SNPs with the most significant sdMAF in the phase 3 data

For the eight selected SNPs, two from each of the four regions (NPR, PAR1, PAR2, and PAR3) with the smallest sdMAF p-values in the combined sample, the population-specific sdMAF p-values remain genome-wide significant (Dataset S1 in the Supplementary Information). Moreover, the directions of the sdMAF are consistent across the five super-populations. For example, for rs201194898 in NPR (GRCh37 position=9,377,082), the sdMAF p-values are <1e-200, 8.82e-149, 1.12e-49, 2.50e-89, 2.91e-45, and 1.95e-101, respectively in the ALL (combined) and the EAS, EUR, AFR, AMR, and SAS super-populations; the corresponding female minus male sdMAFs are all >0.27. That is, the MAFs of rs201194898 are significantly larger in females than in males, across all five super-populations. Note that the minor allele, defined based on sex- and population-combined sample, may not have MAF less than 0.5 in a sex- and/or population-stratified sample, and for a SNP in NPR and PAR3 each male only contributes a single allele to the MAF calculation. Similar results are reported in Dataset S1 for rs6634333 in NPR, as well as the two PAR3 SNPs in the 88,460,295 and 88,462,611 positions (GRCh37).

For the PAR1 SNP at 2,697,599, the super-population sdMAF p-values are all <1e-200 (except 2.69e-80 in AMR); the corresponding female minus male sdMAFs are all more extreme than −0.46 (−0.34 in AMR). Similar results were observed for the PAR2 SNP at 154,934,295, for which sdMAF p-values <1e-200 in all five super-populations: MAFs are significantly smaller in females than males. In addition, all females in EAS, EUR and AMR are homozygous TT, while males in EAS, AFR and AMR are all AT heterozygotes. In other words, genotype is fixed by sex in EAS and AMR.

The population-specific HWE testing results, however, vary across the eight SNPs or across the five super-populations. For example, for rs6634333 in NPR, the HWE testing p-values in females are genome-wide significant in all but SAS: 1.71e-30, 3.21e-34, 1.09e-18, 1.97-22, and 2.41e-4 respectively in EAS, EUR, AFR, AMR and SAS. The corresponding population-specific HWD delta estimates are all negative, with an excess of heterozygous females. For rs201194898 in NPR and the two PAR3 SNPs, population-stratified HWE testing in females are genome-wide significant, with excesses of heterozygous females in all five super-populations.

For the PAR1 SNP in the 2,697,599 position where HWE testing could be performed in both females and males, HWE testing in males are all genome-wide significant, but in females the HWE p-values >0.8; consistent results were observed for the other three SNPs in PAR1 and PAR2 (Dataset S1). In addition, a SNP can be monomorphic in females (HWE p-value = NA in that case), while in the male sample the heterozygous genotype may be present for some of the five super-populations.

### Comparison to autosomal 1, 7 and 22 phase 3 data

Chromosomes 1, 7 and 22 were selected as the longest, most similar in length to the X chromosome, and one of the shortest. There were 530,434, 406,057 and 97,216 biallelic SNPs with global MAF ≥5% on these chromosomes, respectively. Manhattan plots of sdMAF p-values (Figures S4, S5 and S6), as well as the histograms and QQ plots (Figure S7), show that there are very few SNPs on these autosomes with genome-wide significant sdMAF. Specifically only 6, 1 and 0 SNPs on chromosomes 1, 7 and 22 had sdMAF p-values <5e-8, respectively.

To obtain better insight into the nature and source the sdMAF on the autosomes, we then further examined six SNPs, two from each of the three autosomes with the smallest sdMAF p-values (Dataset S1). For example, on chromosome 1 rs10803097 (GRCh37 position=243050350; sdMAF p-value =2.15e-25) has higher MAFs in males than females in all five super-populations, but the population-specific sdMAFs are only genome-wide significant in EAS and AFR (Dataset S1) while sdMAF p-value = 0.12 in SAS, suggesting heterogeneity in sdMAF between the super-populations at this autosomal SNP. The HWE testing are genome-wide significant in males in all super-populations (except SAS) with excess of heterozygous males, but not in females. A BLAST (11) search of a 100 nucleotide sequence flanking this SNP identified perfect match to sequence on the NPR region of the Y chromosome (GRCh38 position:11786038), with the Y chromosome having the alternate allele at the SNP, suggesting that it is a PSV (12).

In contrast, on chr 7 rs78984847 (GRCh37 position=72053830; sdMAF p-value =3.5e-18) has higher MAFs in females than males in all five super-populations. The population-stratified sdMAF p-values are consistently small, 7.66e-5, 3.06e-4, 1.76e-5, 7.52e-3, and 1.79e-6 respectively in EAS, EUR, AFR, AMR and SAS, but not genome-wide significant as a result of reduced sample sizes. In addition, HWE p-values are much smaller in females than males for all five super-populations, with excess of heterozygous females; the HWE p-values in females are <5e-8 in EUR, AFR and SAS, and 2.92e-7 in EAS and 1.48e-5 in SAS (Dataset S1). There is also evidence for departure from HWE in males; the HWE p-values in males are 2.02e-2, 1.67e-3, 1.21e-4, 2.57e-4 and 1.57e-2 respectively in EAS, EUR, AFR, AMR, and SAS, with excess of CC males. BLAST of a 100 nucleotide sequenced centred on this SNP identified multiple close matches to other chromosomes, including the X chromosome.

### Comparison between the phase 3 and high coverage sequence data of the X chromosome

Phase-specific sdMAFs and sex-specific genotype agreements between the two phases were examined for 32 SNPs, selected based on the smallest sdMAF p-values in the phase 3 data (GRCh37) from the four regions (NPR, PAR1, PAR2, and PAR3), and success of liftover onto the high coverage data (GRCh38): 10 in PAR1, 4 in NPR, 9 in PAR3, and 9 in PAR2 (Note S2 in the Supplementary Information with SNPs ordered by the GRCh37 positions.)

For the 10 SNPs in PAR1, the high coverage data did not resolve the genome-wide significant sdMAF observed in the phase 3 data (pages 2-11 of Note S2). In addition, they showed good genotype agreement with the phase 3 data; similar results were observed for 9 SNPs in PAR2 (pages 25-33 of Note S2). For the 4 SNPs in NPR (pages 12-14 and 24 of Note S2), sdMAFs are no longer genome-wide significant in the high coverage data, suggesting genotyping error in phase 3. For the 9 SNPs in PAR3 (pages 15-23 of Note S2), two persisted with genome-wide sdMAF in the high coverage data, located at the centromeric boundary between PAR3 and NPR, while the remaining seven in PAR3 were no longer significant.

Results of X chromosome-wide sdMAF analysis of the high coverage data, without the liftover restriction, are reported in Table S2 and Figures S8-S14 of the Supplementary Information.

Figure S8 shows the analytical pipeline of selecting bi-allelic SNPs with population-pooled MAF ≥5% using the high coverage data, and Table S2 contains the counts of variants by region, MAF threshold and those excluded.

The Manhattan plot of sdMAF p-values in Figure S9 shows that, as compared to Figure 2 for the phase 3 data, the prevalence of genome-wide significant sdMAF is reduced in the high coverage data for NPR and PAR3, about a 10-fold deduction: 0.11% of SNPs in NPR and 0.07% in PAR3 have sdMAF p-values <5e-8 in the high coverage data, as compared to 0.83% in NPR and 0.85% in PAR3 in phase 3. This suggests that genotyping error is a contributing factor to some of the significant sdMAF observed in the phase 3 data. The causes of the remaining sdMAF in NPR and PAR3 in the high coverage data require further examination.

The sdMAF results for PAR1 (Figures 3 and S10) and PAR2 (Figures 4 and S11) are practically the same between the two phases: 0.30% of SNPs in PAR1 and 12.2% in PAR2 have sdMAF p-values <5e-8 in the high coverage data, as compared to 0.29% in PAR1 and 13.1% in PAR2 in phase 3. The sdMAF SNPs in the high coverage data still occur at the NPR-PAR1 and NPR-PAR2 boundaries, which are evident from the zoomed-in Manhattan plots (Figures S10 and S11)..

SNPs with significant sdMAF also remain at the centromeric NPR-PAR3 boundary (Figure S12). The persistent presence of small sdMAF p-values, including in the NPR region, is also evident from the QQ plots (Figure S13) and histograms (Figure S14). This strongly suggests that sex-linkage is a major driver for sdMAF at both PAR1 and PAR2, and at the centromeric boundary of PAR3. Association studies of the X chromosome thus must consider sdMAF.

## Discussion

Our initial sdMAF analysis focused on the phase 3 data since it has been examined extensively for association analysis, and is one of the most commonly used imputation panels for GWAS (13). With the recent release of the high coverage data, we first compared results between the two phases for specific SNPs. In addition, we also did a separate X chromosome-wide analysis of the high coverage data since 58% of phase 3 X chromosomal SNPs could not be lifted over from GRCh37 to GRCh38 (14).

We identify two likely sources of sdMAF: genotyping error and sex-linkage. Genotyping error accounts for many NPR and PAR3 sdMAF in phase 3, since they were mostly resolved in the high coverage data. However, sdMAF NPR and PAR3 SNPs remain in the high coverage data. Thus, despite the recent advance in how to analyze NPR and PAR3 SNPs (6), our findings here show that robust association methods for analyzing the X chromosome must consider sdMAF caused by genotyping error.

In contrast, the impact of sex-linkage in PAR1 and PAR2 results in most sdMAF persisting in the high coverage data. Sex-linkage at PARs has previously been discussed for linkage analyses using affected sibpairs (7), but it has not been examined in the context of association studies (15, 16). Multiple authors have stated that association methods routinely used for autosomes can be applied to the PARs (2, 4, 5). However, our results indicate that association analyses at the PAR-NPR boundaries (and at the centromeric boundary of PAR3) should consider sdMAF caused by sex-linkage.

We searched the NHGRI-EBI GWAS catalog (17) and identified multiple signals at the PAR boundaries. These include loci: for BMI and multiple lipids at the PAR2 boundary (18, 19); ANCA-associated vasculitis at PAR3 (20, 21); age of onset of myopia close to the PAR1 boundary (21); and a separate PAR1 locus for mean corpuscular volume (22). None of these studies provided sex-specific results, making it difficult to determine the effect of sdMAF on these associations.

Of note, for many variants with sdMAF, there was also deviation from Hardy-Weinberg equilibrium in each of the super-populations in females (NPR and PAR3) and males (PAR1 and PAR2) (Dataset S1). Hardy-Weinberg equilibrium testing of variants in the 1000 Genomes phase 3 data has been examined separately in the JPT and YRI populations (23). SNPs with missing rate <5% and possessing an rs identifier were used. The earlier work (23) found lower rates of deviation from HWE on the X chromosome (after exclusion of PAR1 and PAR2) than on any of the autosomes, but the sample size for the X chromosome is ∼3/4 of that for the autosomes. Results from our sdMAF analysis also calls for new X chromosome-aware HWE methods that consider sdMAF.

### Online Methods

#### The X chromosome phase 3 data of the 1000 Genomes Project

The 1000 Genomes Project generated an integrated call set of variants for phase 3 (release 5) data based on four data types: Illumina 2.5M genotyping array, Affymetrix SNP6.0, high-coverage whole exome sequence (WES), and low coverage whole genome sequence (WGS) (24). Because only a minority of variant positions are covered by the first three, >93% of the data come from the WGS, which is generally of low quality due to its low-coverage nature (average depth of 7.4 on the autosomes).

The phase 3 data of the 1000 Genomes Project identified 3,468,093 variants on the X chromosome (24), with an average depth of 6.2. Assuming equal proportions of males and females, the sex-specific average depth is 7.0 for females and 3.5 for males.

Focusing on bi-allelic common SNPs presumed to be of high quality, we first removed variants that had >2 alleles, as well as indels. We then analysed all the bi-allelic SNPs from the whole of the X chromosome, including the pseudo-autosomal region 1, PAR2 and PAR3 (8), in addition to the non-pseudo-autosomal region. X chromosomal locations for these four regions, NPR, PAR1, PAR2, and PAR3, were obtained from The Genome Reference Consortium, based on GRCh37.

SNPs were included based on global minor allele frequency ≥5%, because they have been shown to have the highest quality in this data (24). We defined the minor allele based on the overall sample of the1000 Genomes Project, with males providing a single allele count in the non-PAR regions for the MAF calculation. The definition of a minor allele has no necessary relationship to the reference allele in GRCh37, and the minor allele defined in the sex- and population-pooled sample may not be minor in sex- or population-stratified samples. The counts of variants by region, MAF threshold and those excluded are provided in Table S1; the analytic pipeline and flow-chart of the analyses are provided in Figure 1; Supplementary codes for all the analytical steps are also provided (Web Resource).

#### Testing for X chromosomal variants with sex difference in MAF (sdMAF)

Sex difference in MAF is an indication of either potential genotyping error or a biological phenomenon particularly at the NPR-PAR boundaries. Although sdMAF analyses could be performed separately for each of the five super-populations, or each of the 26 populations, this is not a powerful approach. As the sex ratio is similar across the populations (Figure S1), we tested sdMAF using the whole sample of the 1000 Genomes Project. This more powerful approach can detect sex difference in MAF if it is present in any of the sub-samples. If there are no sdMAFs in any of the sub-samples, the test remains valid (i.e. accurate control of false positives) even if the MAFs differ drastically between the sub-populations. This is because the sdMAF test detects the difference in MAF *between females and males*, not the difference in MAF between populations.

#### sdMAF test

For each bi-allelic SNP in the NPR and PAR3 regions of the X chromosome, notations in Table S3 (A) denote sex-stratified genotype counts, where, without loss of generality, allele *A* represents the minor allele defined in the sex-pooled whole sample.

To identify variants in the NPR and PAR3 regions with sex difference in MAF, we used the following conservative test statistic,

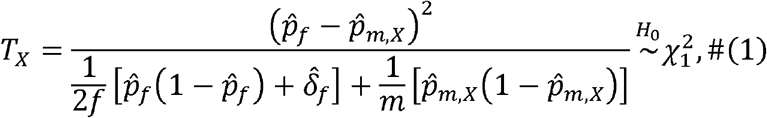

where the numerator,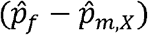, contrasts the frequency estimates of allele *A* between the female and male groups, and the denominator is the estimate of the variance of 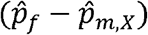 while allowing for Hardy-Weinberg disequilibrium (HWD) in females via 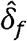. Specifically,

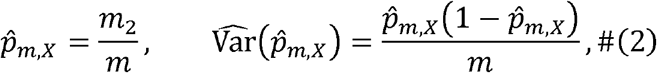

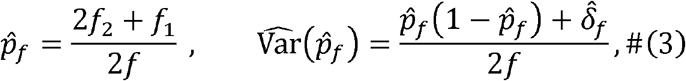

Where

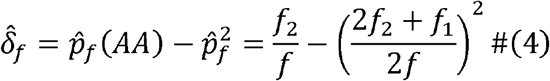

is the estimate of HWD present in females (25). Under the null of no sdMAF, *H*_0_, the test statistic *T*_*X*_ is asymptotically 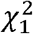 distributed, as it is a straightforward application of the classic two-sample comparison with the consideration of HWD. Note that the *H*_0_ of interest here refers to no sex difference in MAF, while allowing for population difference in MAF.

When applied to a sample consists of individuals from multiple populations (e.g. the whole sample of the 1000 Genomes Project), the sdMAF test based on *T*_*X*_ is conservative because the denominator in equation (1) contains a population-pooled HWD estimate. Note that even if each of the five super-populations or 26 populations is in Hardy-Weinberg equilibrium (HWE), the combined population may not be in HWE unless the MAFs are the same across all populations (26). In addition, we show in the Supplementary Note S1 that the bias factor for the population-pooled HWD estimate is always greater or equal to zero, resulting in *T*_*X*_ being (slightly) conservative in testing for sdMAF.

We note that using a conservative sdMAF test is not an issue for the purpose of this study, because for SNPs declared to have significant sdMAF we are then more confident about the conclusion. Likewise, we used the genome-wide significance level of p-value <5e-8 (27) to declare sdMAF significance, which is conservative for our X chromosome-focused analysis.

Table S3 (B) shows the genotype counts for each bi-allelic SNP in the PAR1 and PAR2 regions, for which a male has three genotypes as for an autosomal SNP. In that case, we used the following test statistic to test for sdMAF,

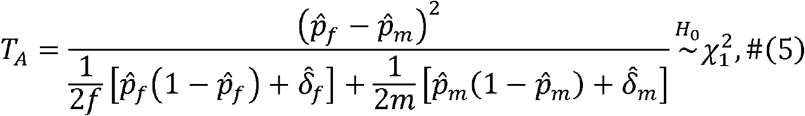

where all notations with subscripts _*f*_ are the same as in equations (3) and (4), while for males,

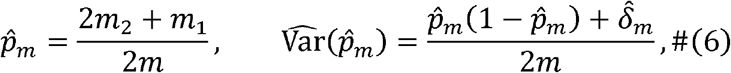

Where

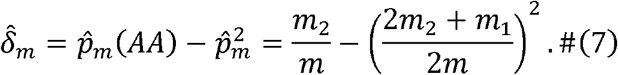

We note that, in finite sample, the Wald’s test shown in equation (5) is more conservative than the Score test, where the sex-stratified MAF and HWD estimates in the denominator of (5) would be replaced with sex-pooled estimates; asymptotically the two tests are equivalent with each other.

#### In-depth analyses of eight X chromosomal SNPs with genome-wide significant sdMAF in the phase 3 data

To better understand the patterns of sdMAF, we selected eight SNPs for additional analyses, two from each of the four regions (NPR and PAR1-3) with the smallest sdMAF p-values in phase 3 data of the 1000 Genomes Project. For each SNP, we calculated population-specific and sex-stratified allele frequency estimates; for a NPR or PAR3 SNP, each male only contributed a single allele count to the allele frequency calculation.

For the four SNPs in the NPR and PAR3 regions, we also performed population-stratified female-only HWE testing for each of the five super-populations, using the standard autosomal method as only females were analyzed here. That is, we used 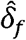 in equation (4) to estimate HWD and the following Pearson’s chi-square test statistic to test for HWD in females,

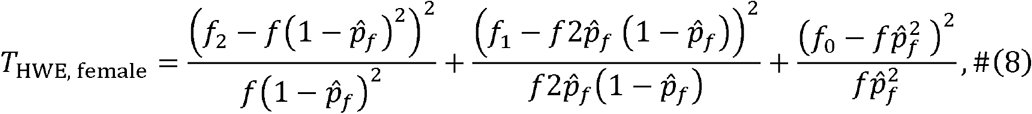

where under the null of HWE, *T*_HWE, femalw_ is asymptotically 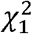 distributed.

We first note that recent work (28) has shown that the Pearson’s chi-square-based HWD test can be reformulated as a test based on the HWD estimate. That is, *T*_HWE, femalw_ in equation (8) can be rewritten equivalently as

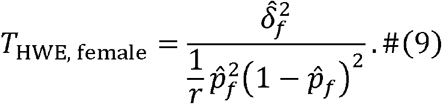

We also note that, although the expressions for 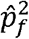 and 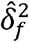 in equation (9) are the same as those in equations (3) and (4), here 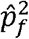 and 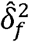 are calculated separately for each of the five super-populations, as HWD testing in the whole sample using the naïve population-pooled HWD estimate is not valid.

For the four SNPs in the PAR1 and PAR2 regions, HWD estimation and testing were also performed in males, as well as jointly with females using sex-pooled estimates. Briefly, we use 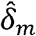 in equation (7) to estimate HWD and use

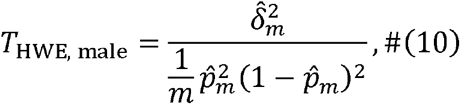

to test for HWD in males, separately for each of the five super-populations. For the sex-combined analysis,

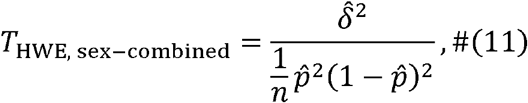

Where

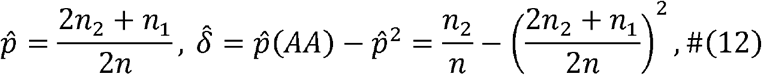

using the notations in Table S3 (B) for a bi-allelic SNP in the PAR1 and PAR2 regions.

#### Analyses of autosomal 1, 7 and 22 phase 3 data for benchmarking

To compare results of variants with significant sex difference in MAF between the X chromosome and autosomes, chromosomes 1, 7, and 22 were first selected to represent the longest, similar size to the X chromosome, and one of the shortest autosomes. Biallelic and common (sex- and population-pooled MAF≥5%) SNPs were then selected for sdMAF analysis, using *T*_*A*_, the sdMAF test statistic shown in (5). When HWE evaluation was warranted, *T*_HWE, femalw_, *T*_HWE, male_ and *T*_HWE,sex - combined_, shown respectively in equations (9), (10) and (11), were applied to the phase 3 data of the 1000 Genomes Project.

#### The X chromosome high-coverage sequence data of the 1000 Genomes Project

To validate the results from the phase 3 data (GRCh37) of the 1000 Genomes Project, we repeated the sdMAF analyses using the recently released high coverage (GRCh38) whole genome sequence data (14).

As noted previously in (14), when phase 3 variants from GRCh37 were lifted onto GRCh38, there was a much higher liftover-failure rate for variants on the X chromosome than for any of the autosomes. Specifically, out of a total of 3,468,093 X chromosomal phase 3 variants (24), 2,020,268 (58%) SNPs and indels failed liftover (14). Therefore, for explicit sdMAF comparison between the phase 3 and high coverage data, we only studied 32 SNPs selected from the four regions of the X chromosome. Specifically, 10, 50, 20, and 50 SNPs, respectively from NPR, PAR1, PAR2, and PAR3, with the smallest sdMAF p-values in the phase 3 data were first selected. Among these SNPs, 10, 4, 9, and 9 SNPs, respectively from NPR, PAR1 and PAR3, were also bi-allelic in the high coverage data and had no missingness in both sets of data. For each of the 32 SNPs, the direction and magnitude of sdMAF were examined, separately for the phase 3 and high coverage data. Genotype agreements between the two sets of data within an individual, separately by sex, were also generated.

Finally, without the liftover constraint between the two phases of the 1000 Genomes Project, we performed an X-chromosome wide sdMAF analysis for the high coverage data using the same sdMAF methods described earlier for the phase 3 data. Supplementary codes for all the analytical steps are also provided (Web Resource).

## Acknowledgements

This research is funded by the Natural Sciences and Engineering Research Council of Canada (NSERC, RGPIN-04934 and RGPAS-522594) and the Canadian Institutes of Health Research (CIHR, MOP-310732).

## Web Resources

The Genome Reference Consortium: https://www.ncbi.nlm.nih.gov/grc/human

The 1000 Genomes Project: https://www.internationalgenome.org

Phase 3 data of the 1000 Genomes Project: http://ftp.1000genomes.ebi.ac.uk/vol1/ftp/release/20130502/ and the specific vcf file used: ftp://ftp.1000genomes.ebi.ac.uk/vol1/ftp/release/20130502/ALL.chrX.phase3_shapeit2_mvncall_integrated_v1b.20130502.genotypes.vcf.gz

High coverage data of the 1000 Genomes Project: http://ftp.1000genomes.ebi.ac.uk/vol1/ftp/data_collections/1000G_2504_high_coverage/working/20190425_NYGC_GATK/ and the specific vcf file used: http://ftp.1000genomes.ebi.ac.uk/vol1/ftp/data_collections/1000G_2504_high_coverage/working/20190425_NYGC_GATK/CCDG_13607_B01_GRM_WGS_2019-02-19_chrX.recalibrated_variants.vcf.gz

## Data and Code Availability

Data used in this work are all publicly available; see Web Resources. All codes used for data analyses are open-resource and available at https://github.com/ZhongWang99

## Supplementary Information

There are three supplementary tables (Tables S1-S3), 14 supplementary figures (Figures S1-S14), two supplementary notes (Notes S1-S2), and one supplementary dataset (Dataset S1) providing additional supporting information.

## Supplementary Information

### Legends of Supplementary Tables

**Table S1. Counts of X chromosome variants by regions, global MAF, and types from the phase 3 data of the 1000 Genomes Project on GRCh37**. See Figure 1 for the analytical pipeline of the SNP selection.

**Table S2. Counts of X chromosome variants by regions, global MAF, and types from the high coverage data of the 1000 Genomes Project on GRCh38**. See Figure S8 for the analytical pipeline of the SNP selection.

**Table S3. Notations of genotype counts for a biallelic SNP on the X chromosome in (A) the NPR and PAR3 regions and (B) the PAR1 and PAR2 regions.** * means not applicable.

### Legends of Supplementary Figures

**Figure S1. Counts of males and females by population in the 1000 Genomes Project phase 3 data on GRCh37**. The populations are first ordered by the 5 super-populations alphabetically, and then by the total counts within each super-population. A: counts; B: the corresponding proportion of males. The red horizontal line represents 0.5.

**Figure S2. QQ plots of the sdMAF p-values of the X chromosome from the 1000 Genomes Project phase 3 data on GRCh37**. Results of bi-allelic SNPs with global MAF ≥5% are shown separately by region, A: NPR; B: PAR1, C: PAR2; D: PAR3. For better visualization p-values < 1e-300 are plotted as 1e-300 (300 on −log10 scale). The red dashed line represents the line of equality. The corresponding Manhattan plots are in Figures 2 (across the whole X chromosome) and Figures 3, 4 and 5 for PAR1, PAR2 and PAR3, respectively.

**Figure S3. Histograms of the sdMAF p-values of the X chromosome from the 1000 Genomes Project phase 3 data on GRCh37**. Results of bi-allelic SNPs with global MAF≥5% are shown separately by region, A: NPR; B: PAR1, C: PAR2; D: PAR3.

**Figure S4. Manhattan plot for testing for sdMAF across chromosome 1 from the 1000 Genomes Project phase 3 data on GRCh37**. A: sdMAF p-values for bi-allelic SNPs with global MAF ≥5% presumed to be of high quality. Y-axis is −log10(sdMAF p-values) and p-values >0.1 are plotted as 0.1 (1 on −log10 scale) for better visualization. The dashed red line represents 5e-8 (7.3 on the −log10 scale). B: Female - Male sdMAF for the same SNPs in part A.

**Figure S5. Manhattan plot for testing for sdMAF across chromosome 7 from the 1000 Genomes Project phase 3 data on GRCh37**. A: sdMAF p-values for bi-allelic SNPs with global MAF ≥5% presumed to be of high quality. Y-axis is −log10(sdMAF p-values) and p-values >0.1 are plotted as 0.1 (1 on −log10 scale) for better visualization. The dashed red line represents 5e-8 (7.3 on the −log10 scale). B: Female - Male sdMAF for the same SNPs in part A.

**Figure S6. Figure S4. Manhattan plot for testing for sdMAF across chromosome 22 from the 1000 Genomes Project phase 3 data on GRCh37**. A: sdMAF p-values for bi-allelic SNPs with global MAF ≥5% presumed to be of high quality. Y-axis is −log10(sdMAF p-values) and p-values >0.1 are plotted as 0.1 (1 on −log10 scale) for better visualization. The dashed red line represents 5e-8 (7.3 on the −log10 scale). B: Female - Male sdMAF for the same SNPs in part A.

**Figure S7. Histograms and QQ plots of the sdMAF p values for chromosomes 1, 7 and 22 from the 1000 Genomes Project phase 3 data on GRCh37**. Results of bi-allelic SNPs with global MAF ≥5% are shown separately by chromosome: A; chromosome 1; B: chromosome 7; C: chromosome 22. Unlike the X chromosome results in Figure S2, there was no truncation of sdMAF p-values at 1e-300 as the smallest sdMAF p-value is around 1e-25 for any of these three autosomes. The red dashed line represents the line of equality.

**Figure S8. Pipeline for selection of X chromosome biallelic SNPs with global MAF** ≥**5%, presumed to be of high quality, from the 1000 Genomes Project high coverage sequence data on GRCh38**. Variants were placed into the NPR, PAR1, PAR2, and PAR3 regions based on positions available from The Genome Reference Consortium and (8). For detailed counts of variant types and global MAF by regions, see Table S2 in the Supplementary Information.

**Figure S9. Manhattan plot for testing for sex difference in MAF across the X chromosome from the 1000 Genomes Project high coverage sequence data on GRCh38**. A: sdMAF p-values for bi-allelic SNPs with global MAF ≥5% presumed to be of high quality. SNPs in the PAR1 and PAR3 regions are plotted in grey, with PAR3 located around 90 Mb. Y-axis is −log10(sdMAF p-values) and p-values >0.1 are plotted as 0.1 (1 on −log10 scale) for better visualization. The dashed red line represents 5e-8 (7.3 on the −log10 scale). B: Female – Male sdMAF for the same SNPs in part A. For Zoomed-in plots for the PAR1, PAR2 and PAR3 regions see Figures S10, S11 and S12, respectively.

**Figure S10. Zoomed-in plot for testing for sex difference in MAF across PAR1 of the X chromosome from the 1000 Genomes Project high coverage sequence data on GRCh38**. A: sdMAF p-values for bi-allelic SNPs with global MAF ≥5% presumed to be of high quality. Y-axis is −log10(sdMAF p-values) and p-values >0.1 are plotted as 0.1 (1 on −log10 scale) for better visualization. The dashed red line represents 5e-8 (7.3 on the −log10 scale). B: Female - Male sdMAF for the same SNPs in part A.

**Figure S11. Zoomed-in plot for testing for sex difference in MAF across PAR2 of the X chromosome from the 1000 Genomes Project high coverage sequence data on GRCh38**. A: sdMAF p-values for bi-allelic SNPs with global MAF ≥5% presumed to be of high quality. Y-axis is −log10(sdMAF p-values) and p-values >0.1 are plotted as 0.1 (1 on −log10 scale) for better visualization. The dashed red line represents 5e-8 (7.3 on the −log10 scale). B: Female - Male sdMAF for the same SNPs in part A.

**Figure S12. Zoomed-in plot for testing for sex difference in MAF across PAR3 of the X chromosome from the 1000 Genomes Project high coverage sequence data on GRCh38**. A: sdMAF p-values for bi-allelic SNPs with global MAF ≥5% presumed to be of high quality. Y-axis is −log10(sdMAF p-values) and p-values >0.1 are plotted as 0.1 (1 on −log10 scale) for better visualization. The dashed red line represents 5e-8 (7.3 on the −log10 scale). B: Female - Male sdMAF for the same SNPs in part A.

**Figure S13. QQ plots of the sdMAF p-values of the X chromosome from the 1000 Genomes Project high coverage sequence data on GRCh38**. Results of bi-allelic SNPs with global MAF ≥5% are shown separately by region, A: NPR; B: PAR1, C: PAR2; D: PAR3. For better visualization p-values < 1e-300 are plotted as 1e-300 (300 on −log10 scale). The red dashed line represents the line of equality. The corresponding Manhattan plots are in Figures S9 (across the whole X chromosome) and Figures S10, S11 and S12 for PAR1, PAR2 and PAR3, respectively.

**Figure S14. Histograms of the sdMAF p-values of the X chromosome from the 1000 Genomes Project high coverage sequence data on GRCh38**. Results of bi-allelic SNPs with global MAF ≥5% are shown separately by region, A: NPR; B: PAR1, C: PAR2; D: PAR3. The corresponding Manhattan plots are in Figures S9 (across the whole X chromosome) and Figures S10, S11 and S12 for PAR1, PAR2 and PAR3, respectively.

### Legends for Supplementary Notes

**Note S1. Bias in (naïve) population-pooled sample estimate of Hardy-Weinberg disequilibrium (HWD) across multiple populations**.

**Note S2. Comparison of genotypes between phase 3 (GRCh37) and high coverage sequence data (GRCh38) from the 1000 Genomes Project for 23 SNPs selected from the X chromosome**.

In total, 10, 50, 20, and 50 SNPs, respectively from NPR, PAR1, PAR2, and PAR3, with the smallest sdMAF p-values in the phase 3 data were first selected.

Among these SNPs, 10, 4, 9, and 9 SNPs, respectively from NPR, PAR1, PAR2, and PAR3, were also bi-allelic in the high coverage data and had no missingness in both sets of data.

Each page represents the results for one SNP, and SNPs are ordered by the GRCh37 positions.

Within each page, the position of the SNP in phase 3 (build GRCh37) and high coverage (GRCh38) are first provided. Next is the female - male sdMAF difference and the sdMAF p-value. The REF and ALT alleles are also provided for each build. Finally, the counts of the agreement of the genotype calls between the phase 3 and the high coverage data are provided, separately by sex.

### Legends of Supplementary Datasets

**Dataset S1. The 1000 Genomes Project phase 3 data and in-depth analysis results for the eight SNPs selected from the X chromosome and six SNPs selected from the autosomes**. For each of the four regions (NPR, PAR1, PAR2, and PAR3) of the X chromosome and for each of the three autosomes analyzed (chromosomes 1, 7 and 22), two SNPs with the smallest sdMAF p-values in the 1000 Genomes Project population-combined sample were selected. SNPs are ordered based on GRCh37 position.

For each SNP, there are 18 rows, showing the genotype counts and other information for female (F), male (M) and sex-combined (Both) sub-samples for the sex- and population-combined sample (ALL), and the EAS, EUR, AFR, AMR, and SAS super-populations. Thus, there are 18*6=108 rows for the six autosomal SNPs, followed by 18*8=144 rows for the eight X chromosomal SNPs.

Column A: row index 1-252.

Column B = SNP: rs name or. if not available from GRCh37

Column C = CHR: 1, 7, 22, or X

Column D = POS: GRCh37 base pair position

Column E = REGION: NA for an autosomal SNP, and NPR, PAR1, PAR2, or PAR3; SNPs are ordered based on GRCh37 position.

Column F = A1: the A1 allele defined by PLINK; may not be the minor allele in the sex- and population-pooled ALL sample.

Column G = A2: the A2 allele.

Column H = Superpopulation: ALL (the sex- and population-combined sample) and the EAS, EUR, AFR, AMR, and SAS super-populations

Column I = Sex: Both (sex-combined), F (female) or M (male)

Column J = A1A1 count: genotype count of homozygous A1A1; 0 means zero counts; NA means not applicable for certain cells. For a NPR or PAR3 SNP, the M counts of A1A2 are NA as a result of no heterozygous males, and the sex-combined Both counts of A1A1 and A2A2 are also NA due to the X-inactivation uncertainty.

Column K = A1A2 count: genotype count of heterozygous A1A2;

Column L = A2A2 count: genotype count of homozygous A2A2

Column M = A1 count: allele count of allele A1; for a SNP in NPR and PAR3 each male only contributes a single allele count

Column N = A2 count: allele count of allele A2; ; for a SNP in NPR and PAR3 each male only contributes a single allele count

Column O = AF of A1: allele frequency of A1 Column P = AF of A2: allele frequency of A2

Column Q = MA: the minor allele defined based on the sex- and population-pooled ALL sample; NA in other cells.

Column R = HWD.delta: the estimate of delta, the measure of HWD which is freq(A1A1) - freq(A1)* freq(A1); NA for population-pooled ALL sample or the male sample when analyzing a SNP in NPR and PAR3.

Column S = HWE.p: p-values of HWE testing; NA if HWD.delta is NA or the SNP is monomorphic in that sample.

Column T = sdMAF: female - male sex difference in MAF, where the minor allele is defined based on the sex- and population-pooled ALL sample.

Column U = sdMAF.p: p-values of sdMAF testing.

## Notes

### Competing Interest Statement

The authors have declared no competing interest.

