## Supplementary material for "Major sex differences in allele frequencies for X chromosome variants in the 1000 Genomes Project data": Table S1-S3, Figure S1-S14

| | | Biallelic SNP | Multi-allelic SNP | Biallelic INDEL | Multi-allelic INDEL | All |
|---|---|---|---|---|---|---|
| NPR | 1%~5% | 197,011 | 2,128 | 18,074 | 3,331 | 220,544 |
| | >=5% | 222,872 | 2,319 | 32,901 | 6,613 | 264,705 |
| | Total | 3,046,734 | 13,972 | 201,692 | 14,667 | 3,277,065 |
| PAR1 | 1%~5% | 7,899 | 146 | 915 | 120 | 9,080 |
| | >=5% | 13,244 | 214 | 2,033 | 418 | 15,909 |
| | Total | 93,374 | 676 | 5,973 | 692 | 100,715 |
| PAR2 | 1%~5% | 484 | 4 | 55 | 3 | 546 |
| | >=5% | 634 | 5 | 87 | 19 | 745 |
| | Total | 9,348 | 30 | 437 | 34 | 9,849 |
| PAR3 | 1%~5% | 7,242 | 87 | 623 | 97 | 8,049 |
| | >=5% | 9,075 | 101 | 1,076 | 225 | 10,477 |
| | Total | 74,471 | 377 | 5,200 | 416 | 80,464 |
| All | | 3,223,927 | 15,055 | 213,302 | 15,809 | 3,468,093 |

**Table S1. Counts of X chromosome variants by regions, global MAF, and types from the phase 3 data of the 1000 Genomes Project on GRCh37.** See Figure 1 for the analytical pipeline of the SNP selection.

| | | Biallelic SNP | Multi-allelic SNP | Biallelic INDEL | Multi-allelic INDEL | All |
|---|---|---|---|---|---|---|
| NPR | 1%∼5% | 202,421 | 0 | 71,926 | 0 | 274,347 |
| | >=5% | 220,061 | 0 | 68,804 | 0 | 288,865 |
| | Total | 2,232,002 | 0 | 424,960 | 0 | 2,656,962 |
| PAR1 | 1%∼5% | 9,446 | 0 | 4,521 | 0 | 13,967 |
| | >=5% | 13,035 | 0 | 4,889 | 0 | 17,924 |
| | Total | 91,095 | 0 | 27,232 | 0 | 118,327 |
| PAR2 | 1%∼5% | 589 | 0 | 160 | 0 | 749 |
| | >=5% | 663 | 0 | 149 | 0 | 812 |
| | Total | 7,758 | 0 | 978 | 0 | 8,736 |
| PAR3 | 1%∼5% | 7,427 | 0 | 1,968 | 0 | 9,395 |
| | >=5% | 8,757 | 0 | 2,111 | 0 | 10,868 |
| | Total | 63,315 | 0 | 10,844 | 0 | 74,159 |
| All | | 2,394,170 | 0 | 464,014 | 0 | 2,858,184 |

**Table S2. Counts of X chromosome variants by regions, global MAF, and types from the high coverage data of the 1000 Genomes Project on GRCh38.** See Figure S8 for the analytical pipeline of the SNP selection.

| | | | | |
|---|---|---|---|---|
| **A** | | | | |
| Female | $aa$ | $Aa$ | $AA$ | Total |
| | $f_0$ | $f_1$ | $f_2$ | $f$ |
| Male | $a$ | * | $A$ | |
| | $m_0$ | * | $m_2$ | $m$ |
| Total | $n_0$ | $n_1 (= f_1)$ | $n_2$ | $n$ |
| **B** | | | | |
| Female | $aa$ | $Aa$ | $AA$ | |
| | $f_0$ | $f_1$ | $f_2$ | $f$ |
| Male | $aa$ | $Aa$ | $AA$ | |
| | $m_0$ | $m_1$ | $m_2$ | $m$ |
| Total | $n_0$ | $n_1$ | $n_2$ | $n$ |

**Table S3. Notations of genotype counts for a biallelic SNP on the X chromosome in (A) the NPR and PAR3 regions and (B) the PAR1 and PAR2 regions.** * means not applicable.

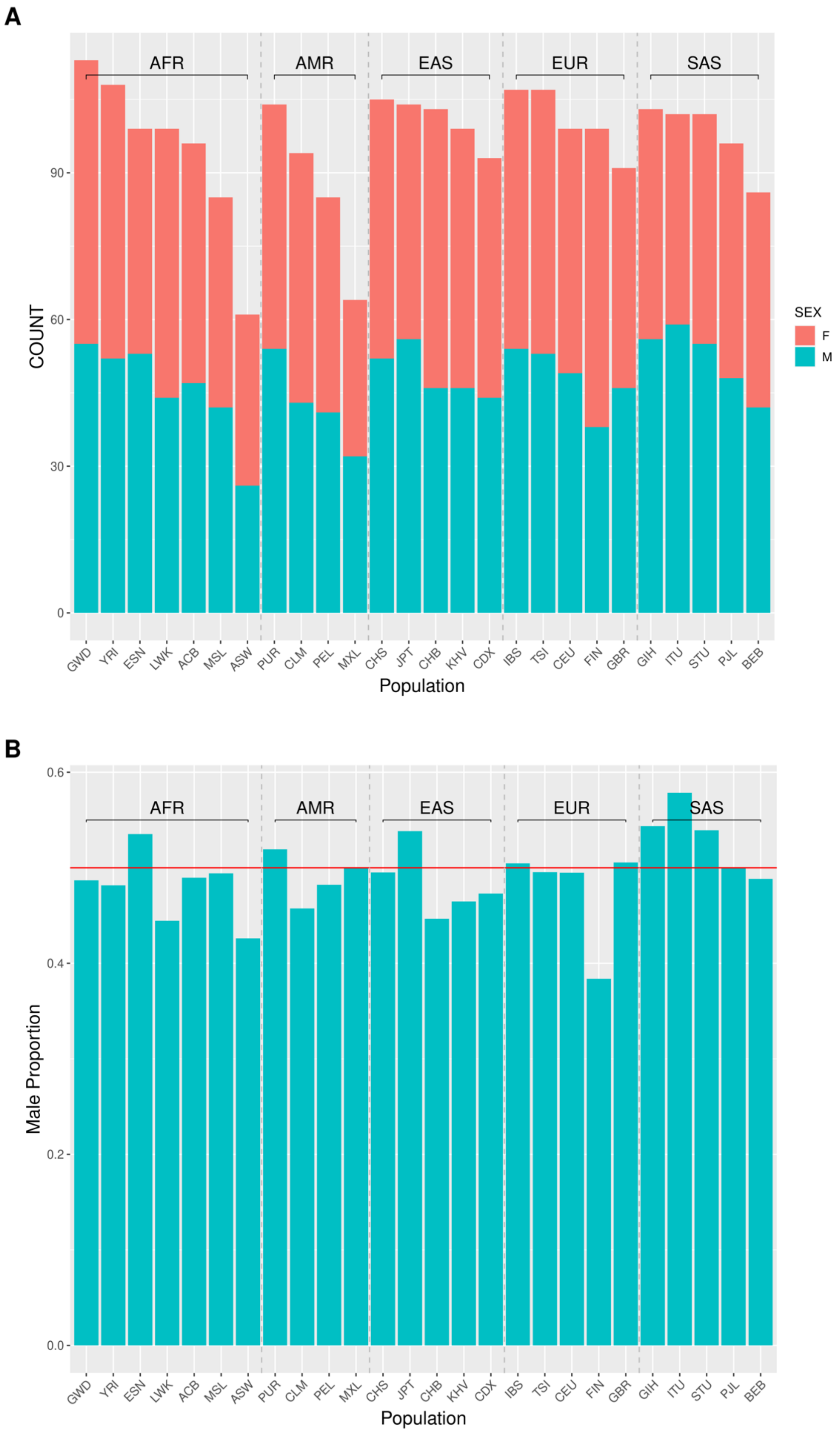
A
AFR
AMR
EAS
EUR
SAS
COUNT
0
30
60
90
SEX
F
M
GWD YRI ESN LWK ACB MSL ASW PUR CLM PEL MXL CHS JPT CHB KHV CDX IBS TSI CEU FIN GBR GIH ITU STU PJL BEB
Population
B
Male Proportion
0.0
0.2
0.4
0.6
AFR
AMR
EAS
EUR
SAS
GWD YRI ESN LWK ACB MSL ASW PUR CLM PEL MXL CHS JPT CHB KHV CDX IBS TSI CEU FIN GBR GIH ITU STU PJL BEB
Population

**Figure S1. Counts of males and females by population in the 1000 Genomes Project phase 3 data on GRCh37**. The populations are first ordered by the 5 super-populations alphabetically, and then by the total counts within each super-population. A: counts; B: the corresponding proportion of males. The red horizontal line represents 0.5.

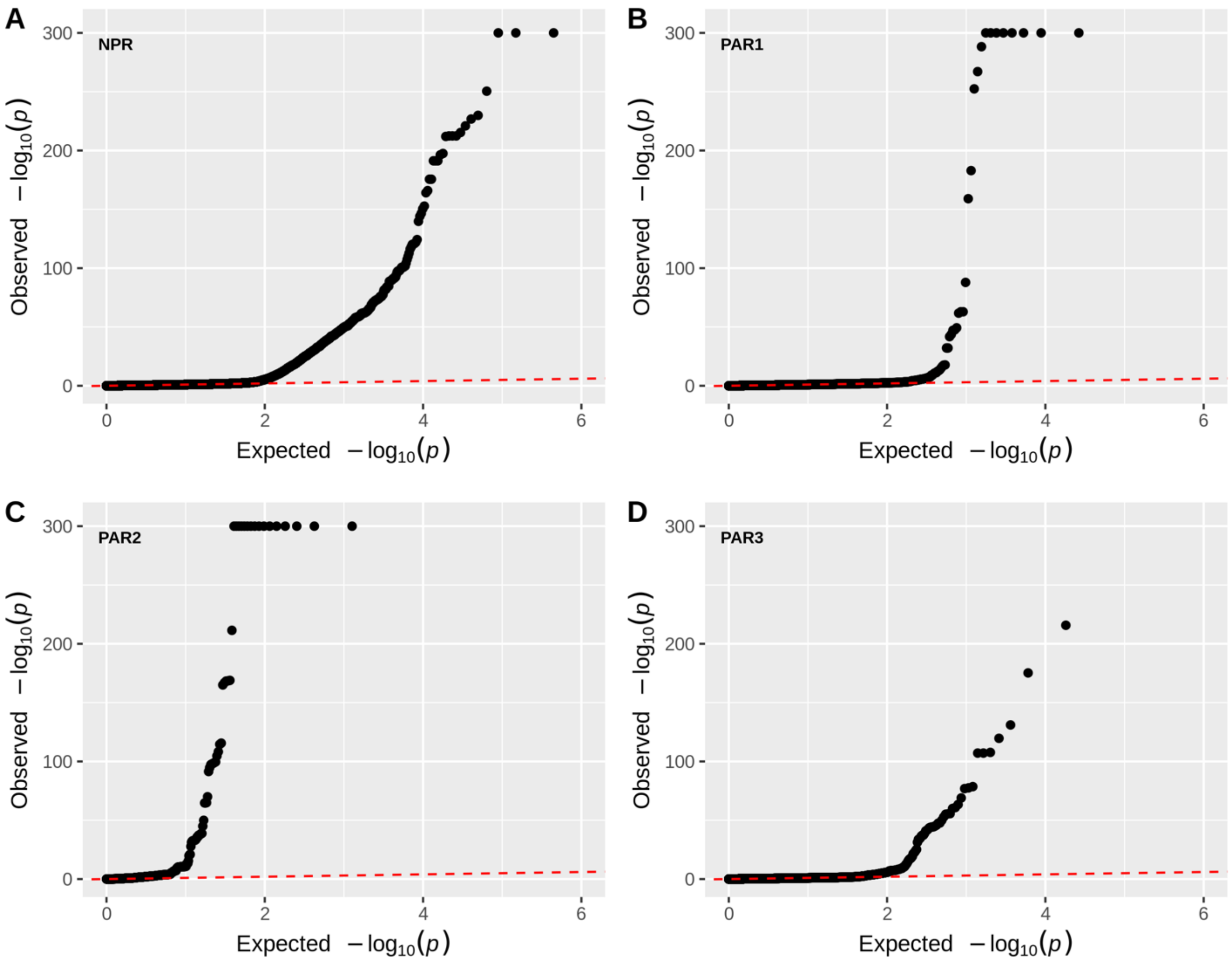


**Figure S2. QQ plots of the sdMAF p-values of the X chromosome from the 1000 Genomes Project phase 3 data on GRCh37.** Results of bi-allelic SNPs with global MAF ≥5% are shown separately by region, A: NPR; B: PAR1, C: PAR2; D: PAR3. For better visualization p-values < 1e-300 are plotted as 1e-300 (300 on -log10 scale). The red dashed line represents the line of equality. The corresponding Manhattan plots are in Figures 2 (across the whole X chromosome) and Figures 3, 4 and 5 for PAR1, PAR2 and PAR3, respectively.

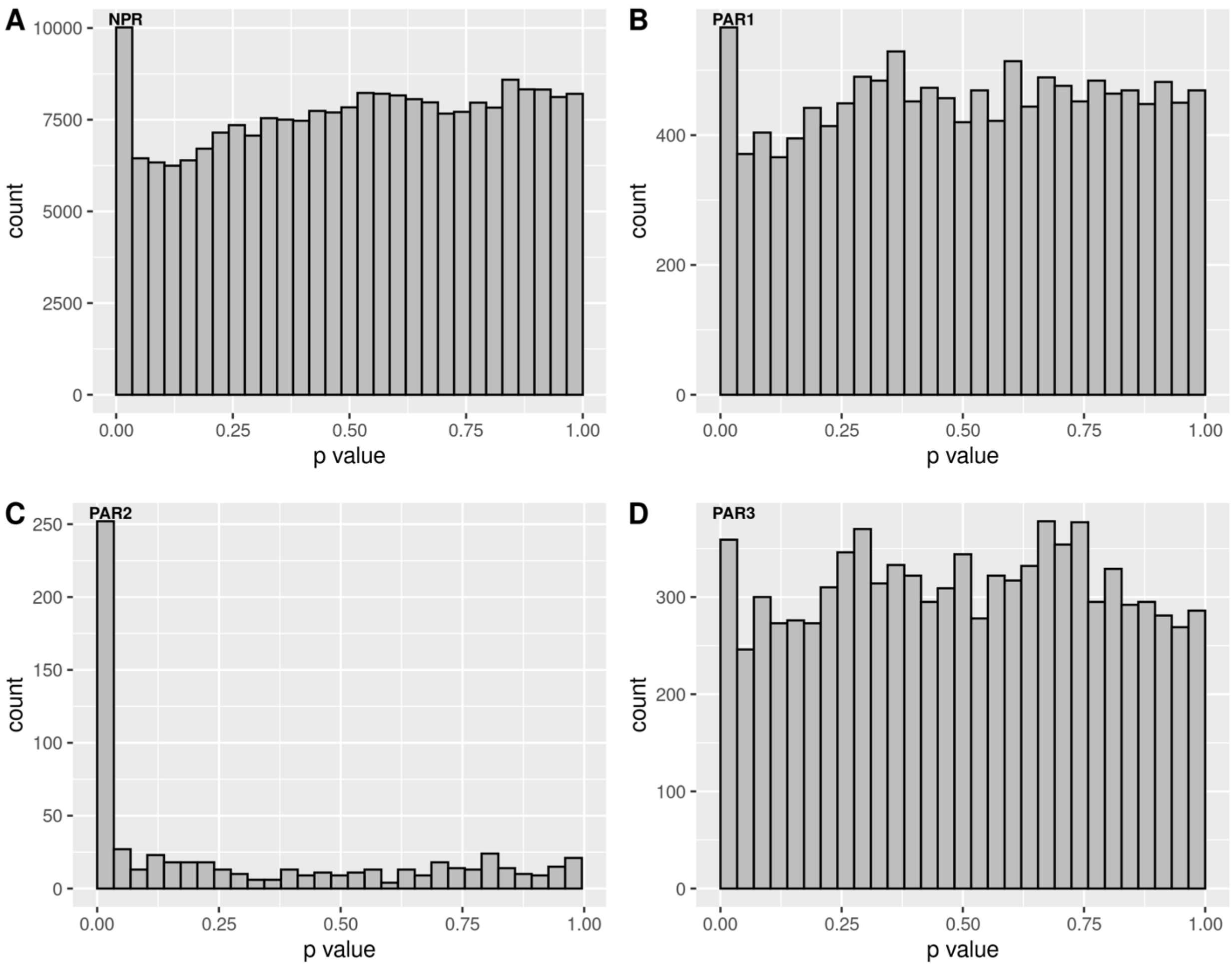


**Figure S3. Histograms of the sdMAF p-values of the X chromosome from the 1000 Genomes Project phase 3 data on GRCh37.** Results of bi-allelic SNPs with global MAF≥5% are shown separately by region, A: NPR; B: PAR1, C: PAR2; D: PAR3.

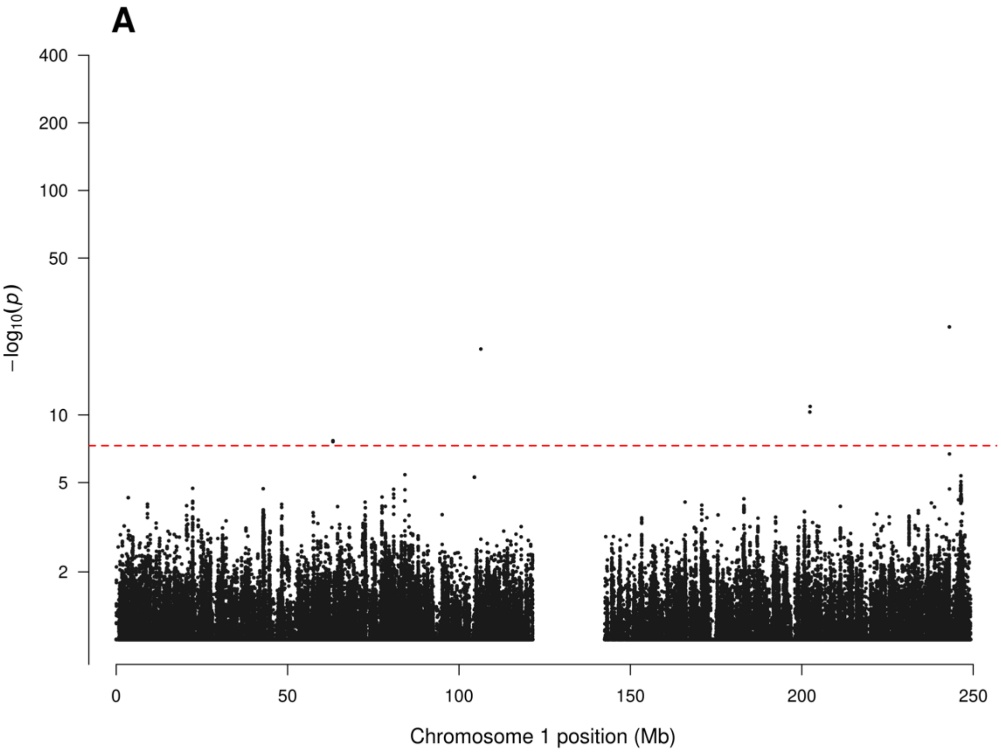

A
400
200
100
50
10
5
2
−log10(p)
0
50
100
150
200
250
Chromosome 1 position (Mb)


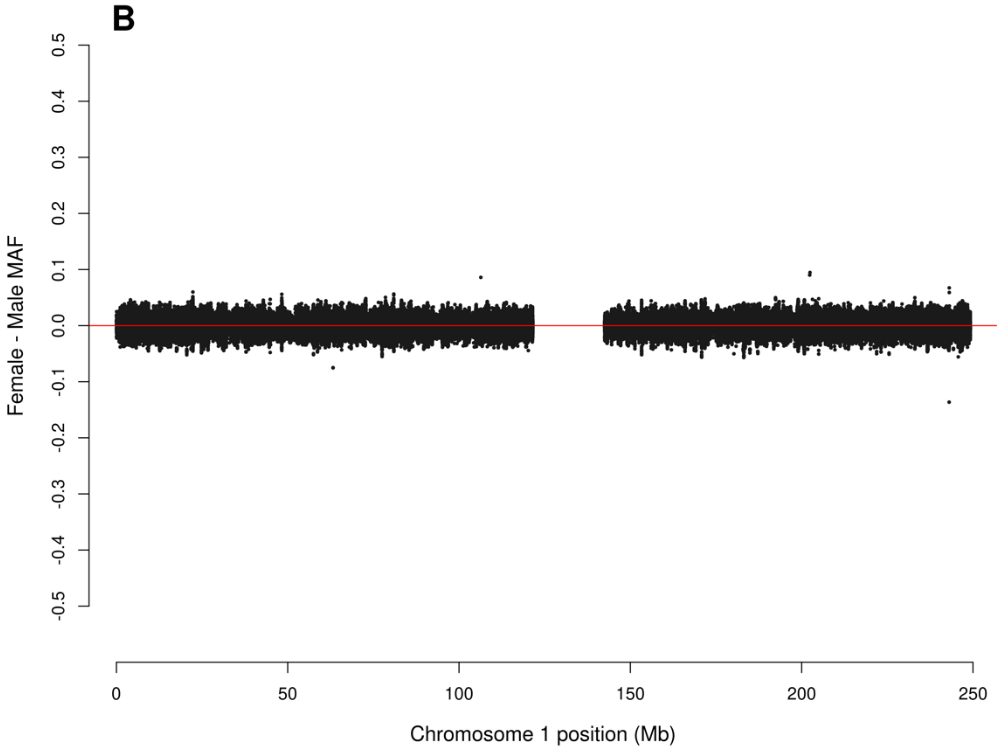

B
0.5
0.4
0.3
0.2
0.1
0.0
-0.1
-0.2
-0.3
-0.4
-0.5
Female - Male MAF
0
50
100
150
200
250
Chromosome 1 position (Mb)

**Figure S4**. **Manhattan plot for testing for sdMAF across chromosome 1 from the 1000 Genomes Project phase 3 data on GRCh37.** A: sdMAF p-values for bi-allelic SNPs with global MAF ≥5% presumed to be of high quality. Y-axis is −log10(sdMAF p-values) and p-values >0.1 are plotted as 0.1 (1 on −log10 scale) for better visualization. The dashed red line represents 5e-8 (7.3 on the −log10 scale). B: Female - Male sdMAF for the same SNPs in part A.

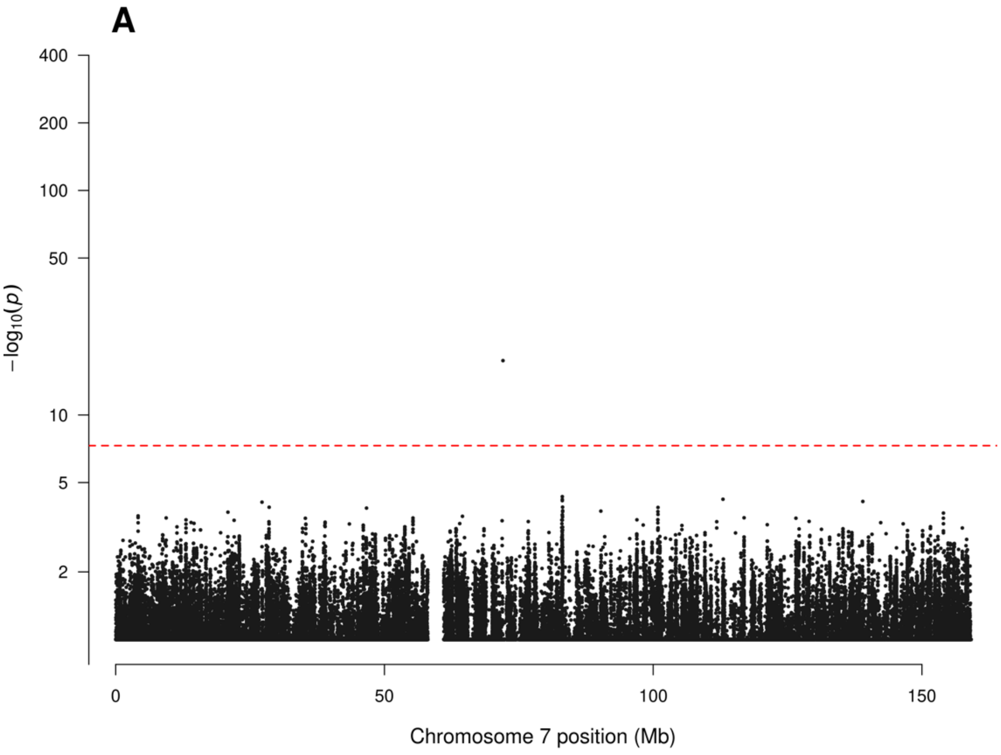
A
400
200
100
50
10
5
2
−log10(p)
0
50
100
150
Chromosome 7 position (Mb)

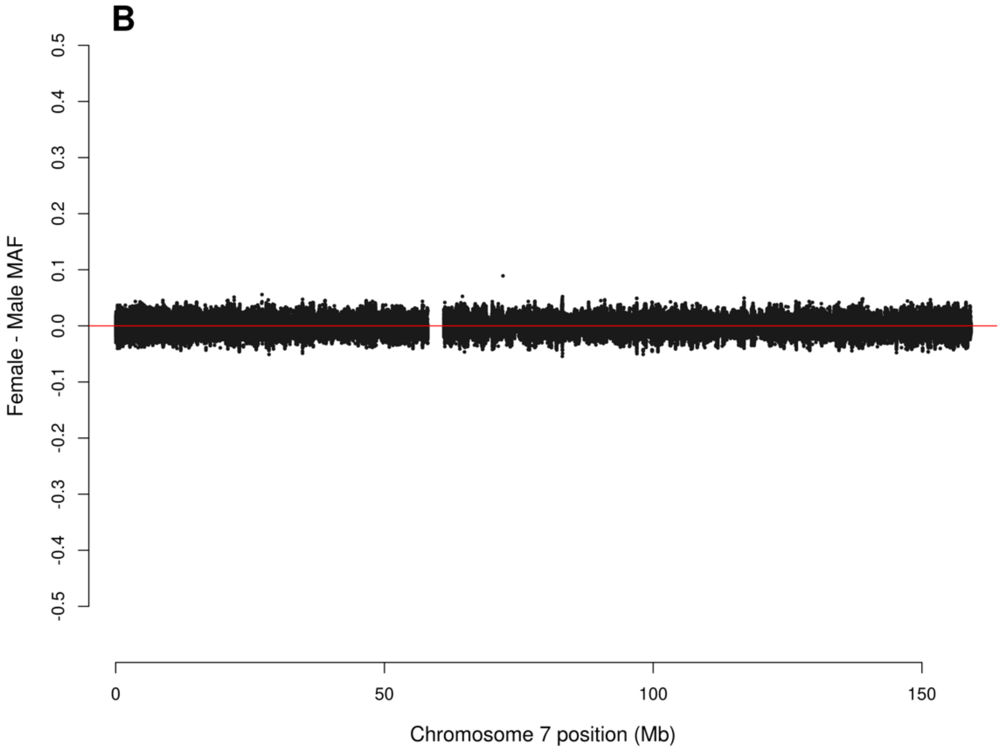
B
0.5
0.4
0.3
0.2
0.1
0.0
-0.1
-0.2
-0.3
-0.4
-0.5
Female - Male MAF
0
50
100
150
Chromosome 7 position (Mb)

**Figure S5**. **Manhattan plot for testing for sdMAF across chromosome 7 from the 1000 Genomes Project phase 3 data on GRCh37.** A: sdMAF p-values for bi-allelic SNPs with global MAF ≥5% presumed to be of high quality. Y-axis is −log10(sdMAF p-values) and p-values >0.1 are plotted as 0.1 (1 on −log10 scale) for better visualization. The dashed red line represents 5e-8 (7.3 on the −log10 scale). B: Female - Male sdMAF for the same SNPs in part A.

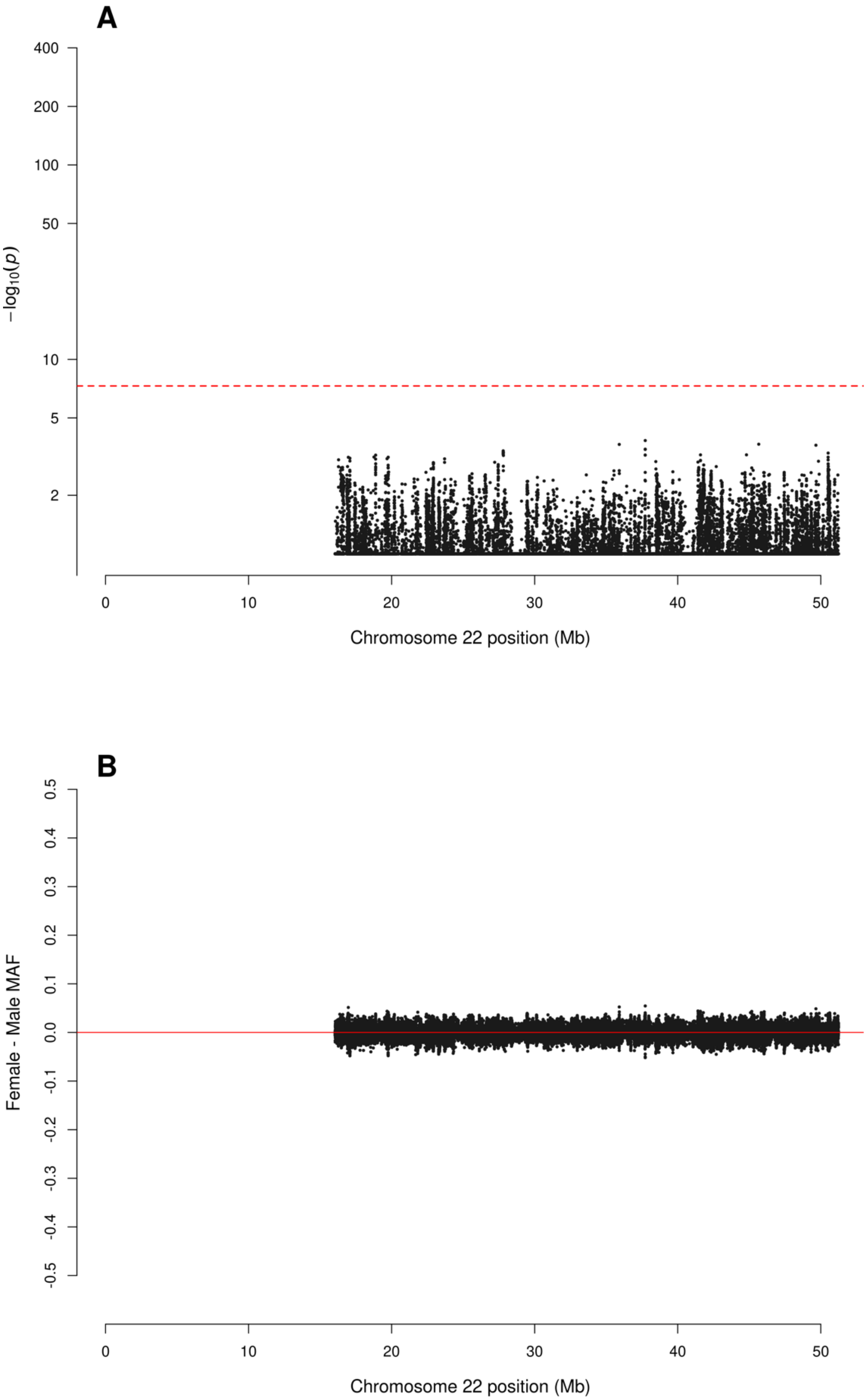
A
400
200
100
50
10
5
2
−log10(p)
0
10
20
30
40
50
Chromosome 22 position (Mb)
B
0.5
0.4
0.3
0.2
0.1
0.0
-0.1
-0.2
-0.3
-0.4
-0.5
Female - Male MAF
0
10
20
30
40
50
Chromosome 22 position (Mb)

**Figure S6**. **Figure S4**. **Manhattan plot for testing for sdMAF across chromosome 22 from the 1000 Genomes Project phase 3 data on GRCh37.** A: sdMAF p-values for bi-allelic SNPs with global MAF ≥5% presumed to be of high quality. Y-axis is −log10(sdMAF p-values) and p-values >0.1 are plotted as 0.1 (1 on −log10 scale) for better visualization. The dashed red line represents 5e-8 (7.3 on the −log10 scale). B: Female - Male sdMAF for the same SNPs in part A.

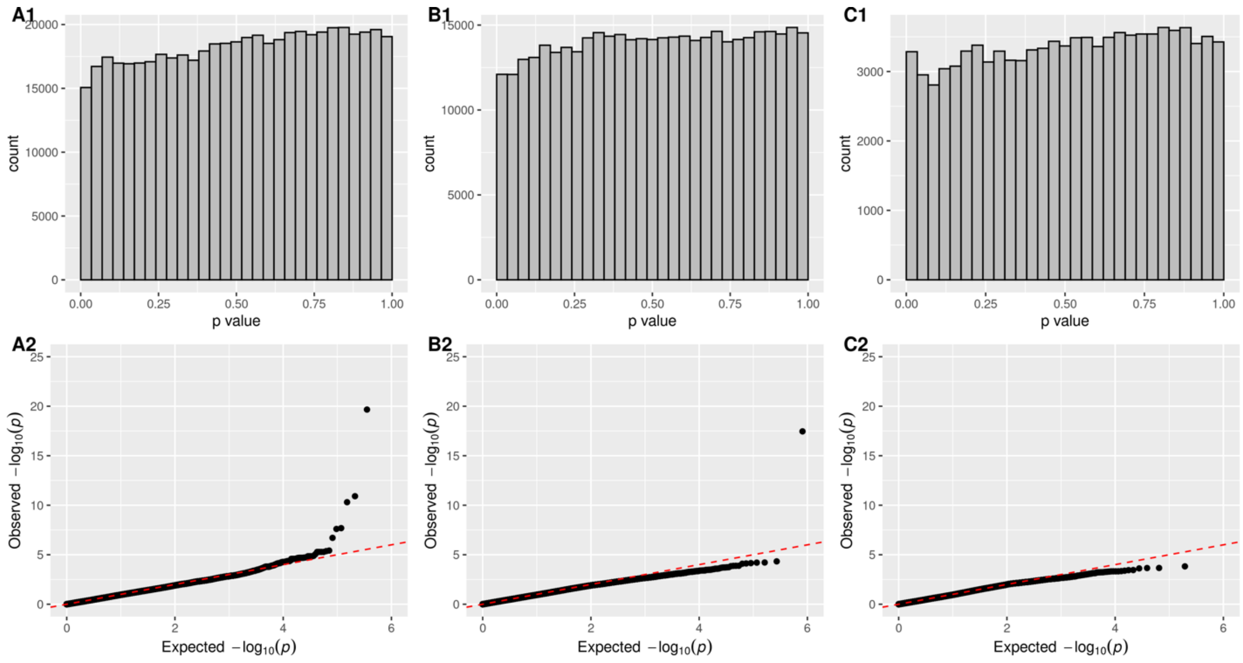


**Figure S7. Histograms and QQ plots of the sdMAF p values for chromosomes 1, 7 and 22 from the 1000 Genomes Project phase 3 data on GRCh37.** Results of bi-allelic SNPs with global MAF ≥5% are shown separately by chromosome: A; chromosome 1; B: chromosome 7; C: chromosome 22. Unlike the X chromosome results in Figure S2 , there was no truncation of sdMAF p-values at 1e-300 as the smallest sdMAF p-value is around 1e-25 for any of these three autosomes. The red dashed line represents the line of equality.

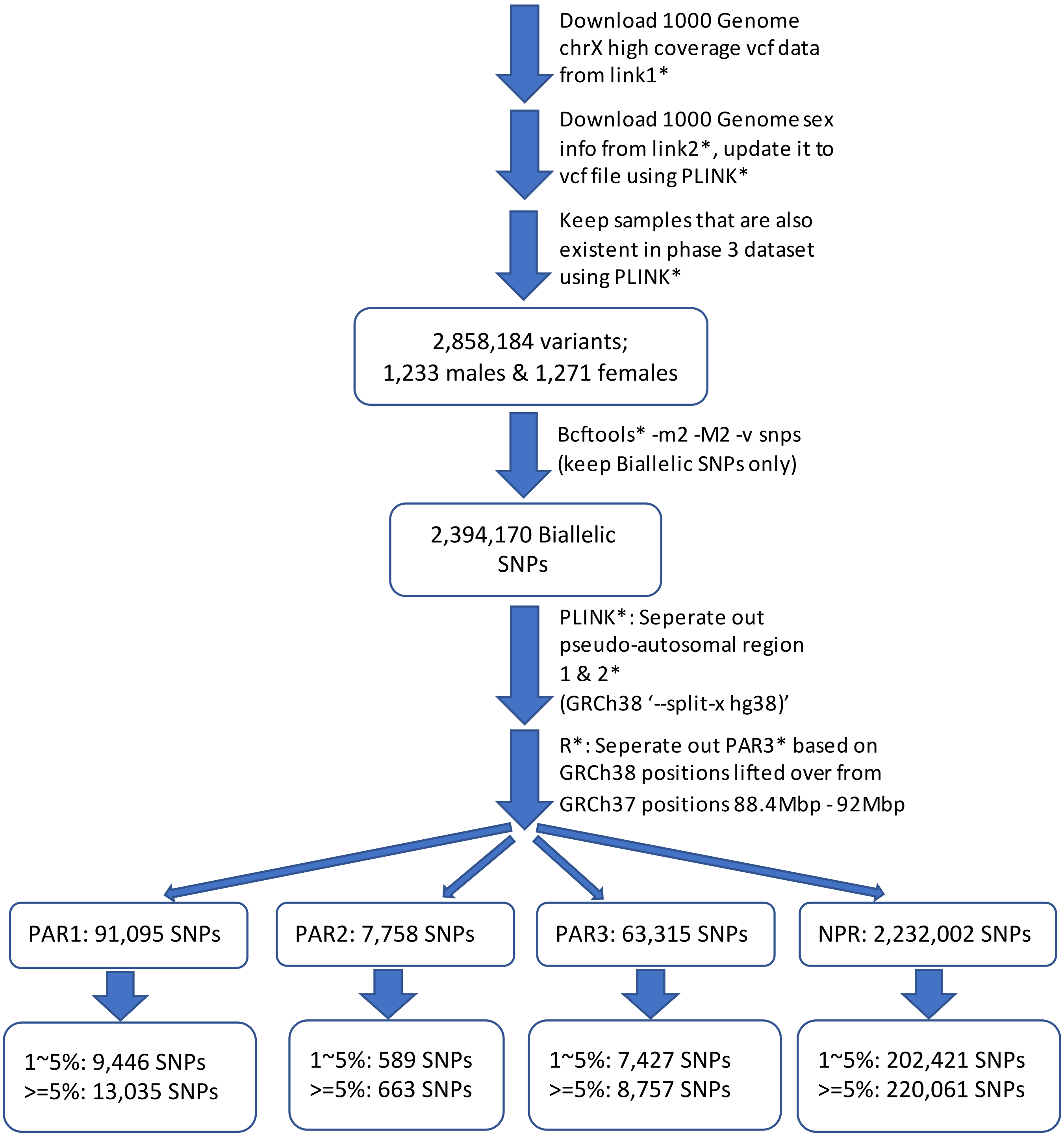


*link1:
http://ftp.1000genomes.ebi.ac.uk/vol1/ftp/data_collections/1000G_2504_high_coverage/working/20201028_3202_phased/CCDG_14151_B01_GRM_WGS_2020-08-05_chrX.filtered.eagle2-phased.v2.vcf.gz
*link2: https://www.internationalgenome.org/data-portal/sample
*PLINK: 1.90 beta version 6.20 64-bit
*PAR1&PAR2 exact location

**Figure S8. Pipeline for selection of X chromosome biallelic SNPs with global MAF ≥5%, presumed to be of high quality, from the 1000 Genomes Project high coverage sequence data on GRCh38.** Variants were placed into the NPR, PAR1, PAR2, and PAR3 regions based on positions available from The Genome Reference Consortium and (8). For detailed counts of variant types and global MAF by regions, see Table S2 in the Supplementary Information.

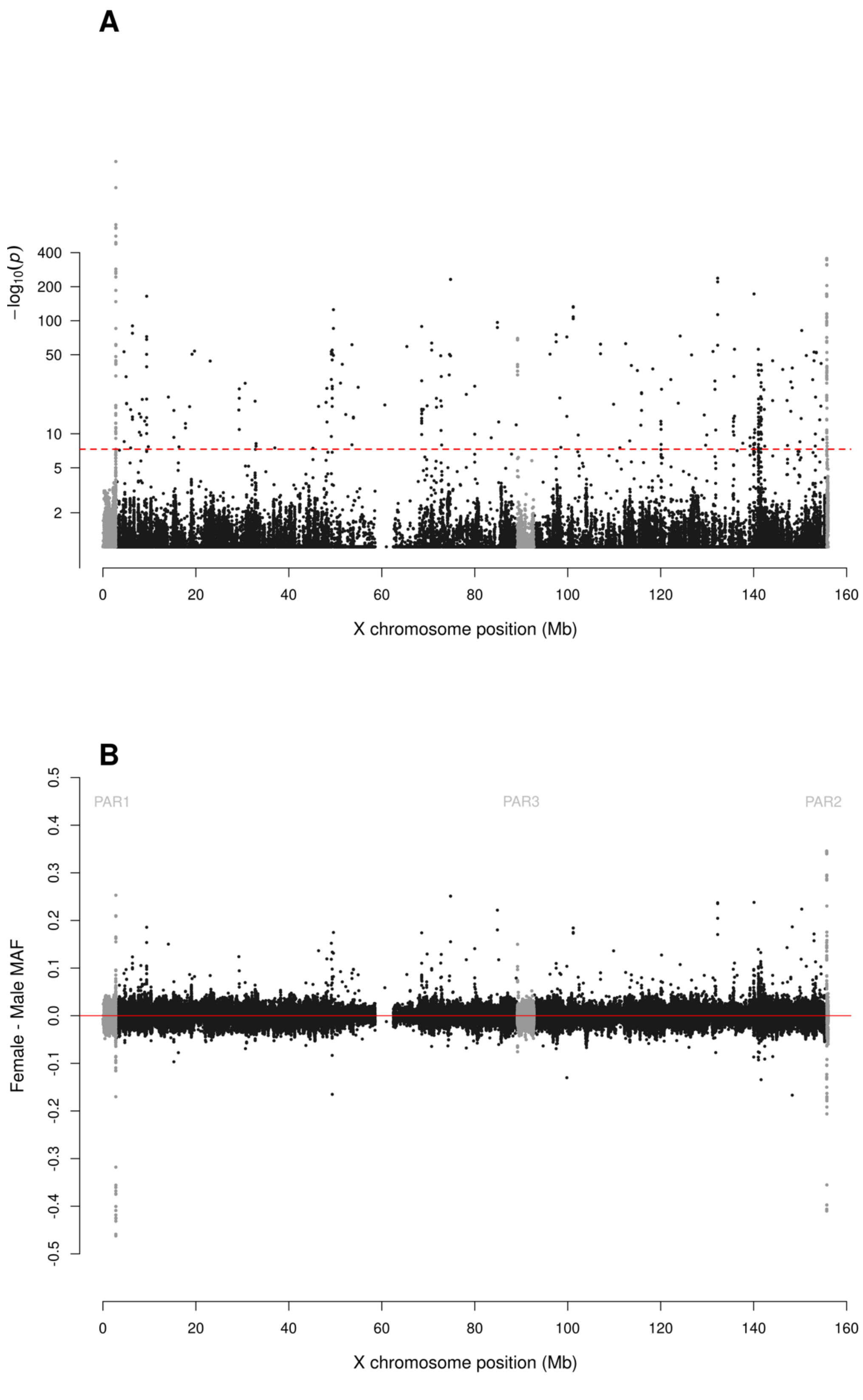
A
$-\log_{10}(p)$
400
200
100
50
10
5
2
0
20
40
60
80
100
120
140
160
X chromosome position (Mb)
B
Female - Male MAF
0.5
0.4
0.3
0.2
0.1
0.0
-0.1
-0.2
-0.3
-0.4
-0.5
PAR1
PAR3
PAR2
0
20
40
60
80
100
120
140
160
X chromosome position (Mb)

**Figure S9. Manhattan plot for testing for sex difference in MAF across the X chromosome from the 1000 Genomes Project high coverage sequence data on GRCh38.** A: sdMAF p-values for bi-allelic SNPs with global MAF ≥5% presumed to be of high quality. SNPs in the PAR1 and PAR3 regions are plotted in grey, with PAR3 located around 90 Mb. Y-axis is −log10(sdMAF p-values) and p-values >0.1 are plotted as 0.1 (1 on −log10 scale) for better visualization. The dashed red line represents 5e-8 (7.3 on the −log10 scale). B: Female - Male sdMAF for the same SNPs in part A. For Zoomed-in plots for the PAR1, PAR2 and PAR3 regions see Figures S10, S11 and S12, respectively.

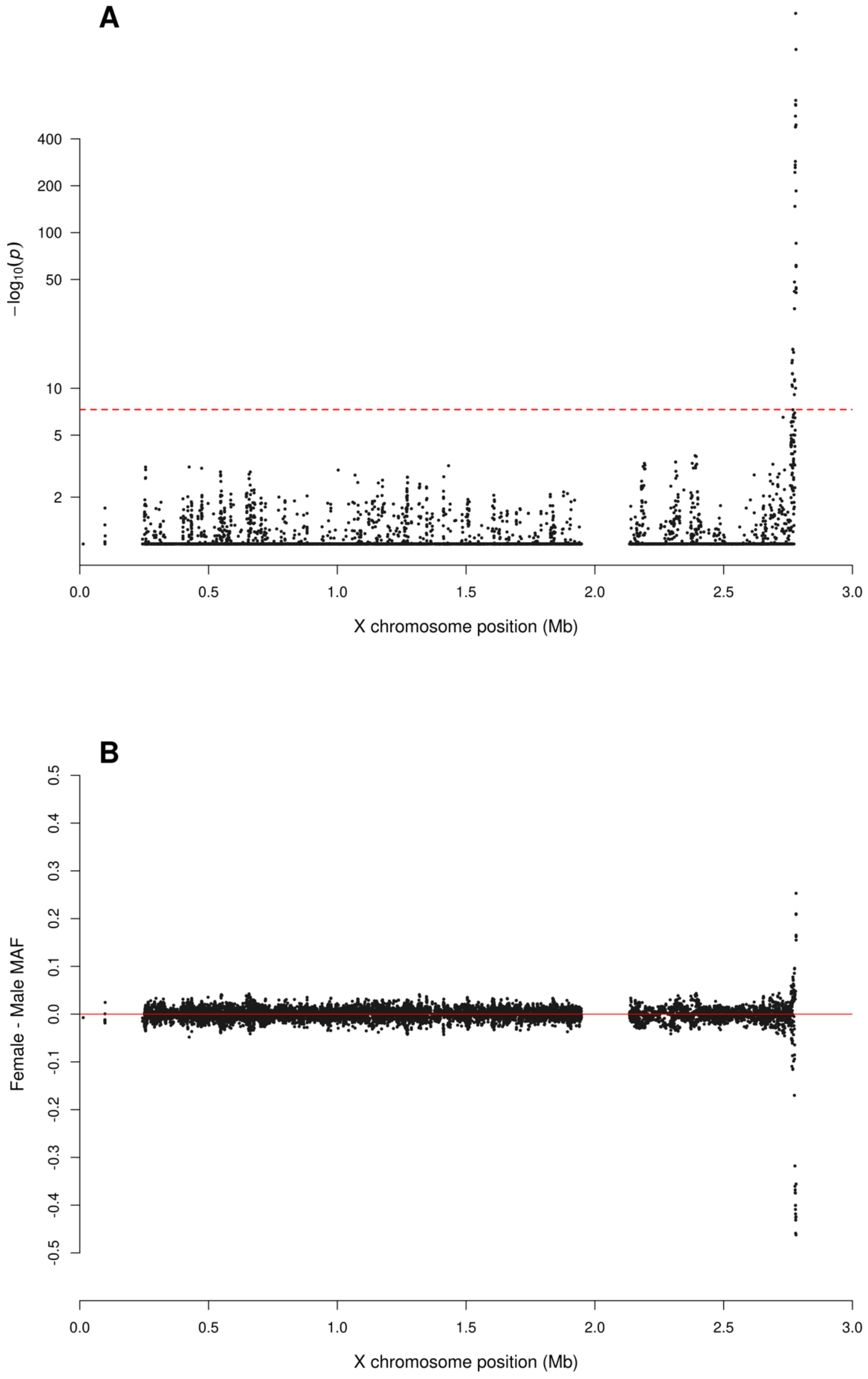
A
−log10(p)
400
200
100
50
10
5
2
0.0
0.5
1.0
1.5
2.0
2.5
3.0
X chromosome position (Mb)
B
Female - Male MAF
0.5
0.4
0.3
0.2
0.1
0.0
-0.1
-0.2
-0.3
-0.4
-0.5
0.0
0.5
1.0
1.5
2.0
2.5
3.0
X chromosome position (Mb)

**Figure S10**. **Zoomed-in plot for testing for sex difference in MAF across PAR1 of the X chromosome from the 1000 Genomes Project high coverage sequence data on GRCh38.** A: sdMAF p-values for bi-allelic SNPs with global MAF ≥5% presumed to be of high quality. Y-axis is −log10(sdMAF p-values) and p-values >0.1 are plotted as 0.1 (1 on −log10 scale) for better visualization. The dashed red line represents 5e-8 (7.3 on the −log10 scale). B: Female - Male sdMAF for the same SNPs in part A.

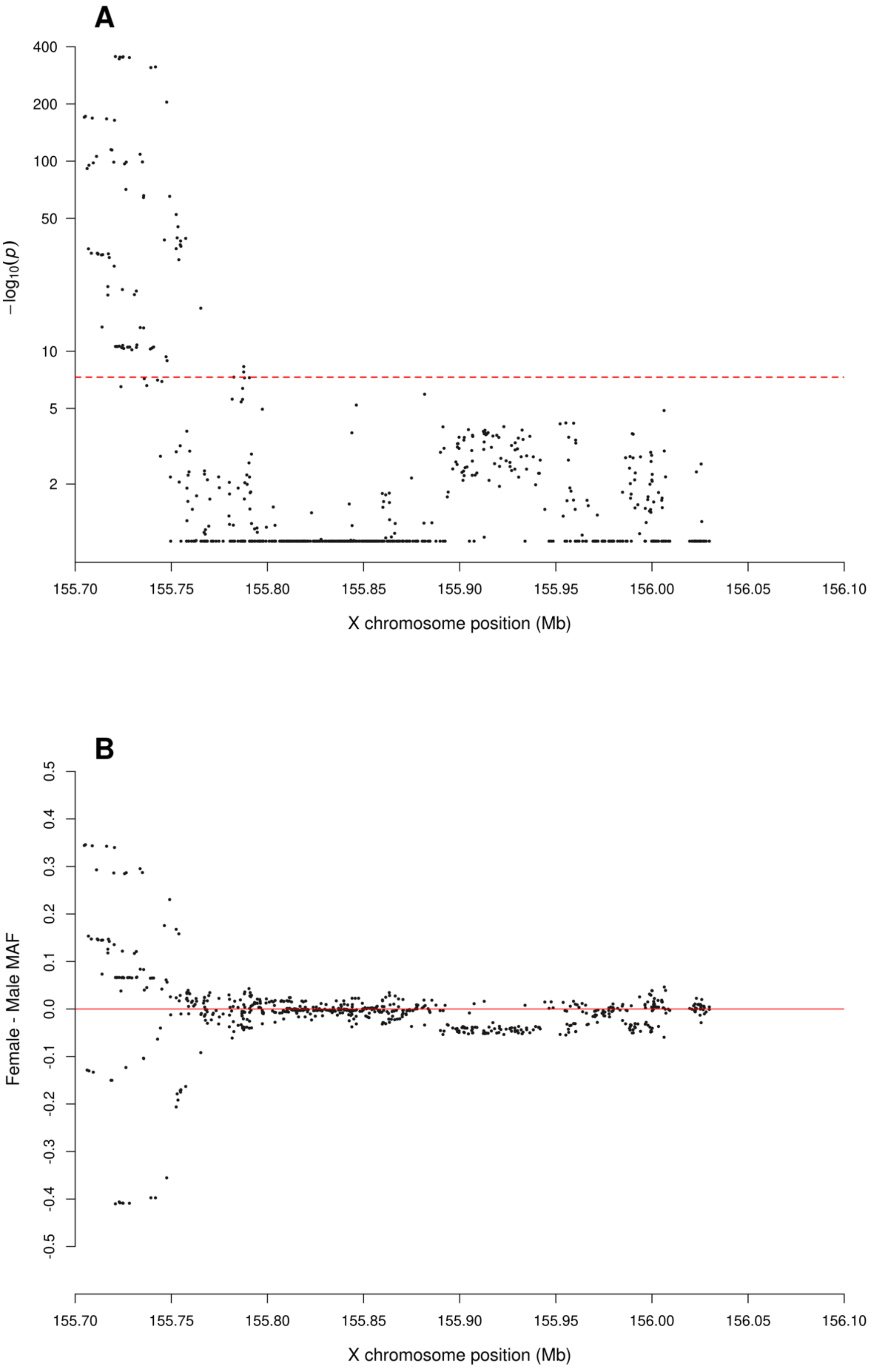
A
400
200
100
50
10
5
2
−log10(p)
155.70
155.75
155.80
155.85
155.90
155.95
156.00
156.05
156.10
X chromosome position (Mb)
B
0.5
0.4
0.3
0.2
0.1
0.0
-0.1
-0.2
-0.3
-0.4
-0.5
Female - Male MAF
155.70
155.75
155.80
155.85
155.90
155.95
156.00
156.05
156.10
X chromosome position (Mb)

**Figure S11**. **Zoomed-in plot for testing for sex difference in MAF across PAR2 of the X chromosome from the 1000 Genomes Project high coverage sequence data on GRCh38.** A: sdMAF p-values for bi-allelic SNPs with global MAF ≥5% presumed to be of high quality. Y-axis is −log10(sdMAF p-values) and p-values >0.1 are plotted as 0.1 (1 on −log10 scale) for better visualization. The dashed red line represents 5e-8 (7.3 on the −log10 scale). B: Female - Male sdMAF for the same SNPs in part A.

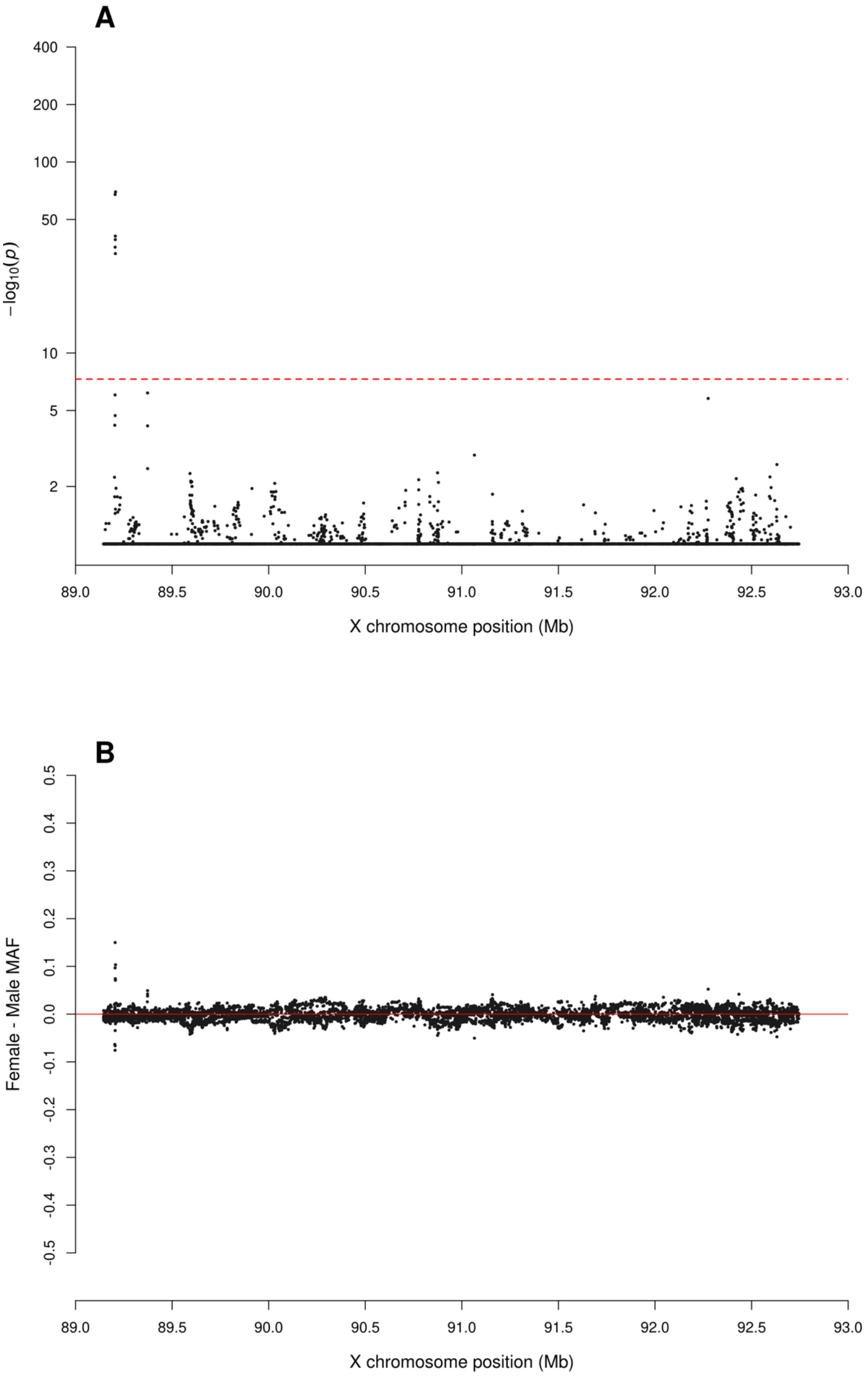
A
400
200
100
50
10
5
2
−log10(p)
89.0
89.5
90.0
90.5
91.0
91.5
92.0
92.5
93.0
X chromosome position (Mb)
B
0.5
0.4
0.3
0.2
0.1
0.0
-0.1
-0.2
-0.3
-0.4
-0.5
Female - Male MAF
89.0
89.5
90.0
90.5
91.0
91.5
92.0
92.5
93.0
X chromosome position (Mb)

**Figure S12**. **Zoomed-in plot for testing for sex difference in MAF across PAR3 of the X chromosome from the 1000 Genomes Project high coverage sequence data on GRCh38.** A: sdMAF p-values for bi-allelic SNPs with global MAF ≥5% presumed to be of high quality. Y-axis is −log10(sdMAF p-values) and p-values >0.1 are plotted as 0.1 (1 on −log10 scale) for better visualization. The dashed red line represents 5e-8 (7.3 on the −log10 scale). B: Female - Male sdMAF for the same SNPs in part A.

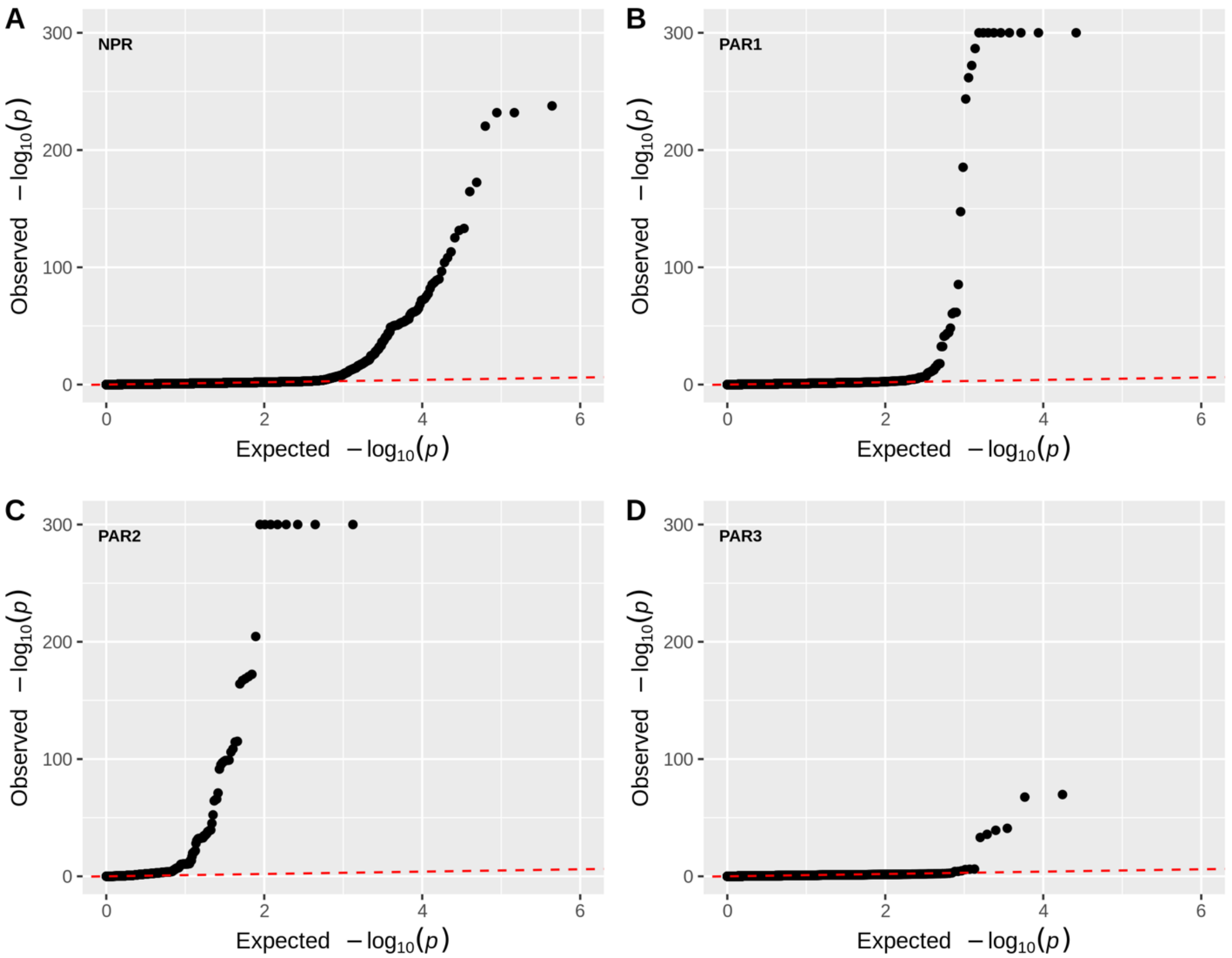


**Figure S13. QQ plots of the sdMAF p-values of the X chromosome from the 1000 Genomes Project high coverage sequence data on GRCh38.** Results of bi-allelic SNPs with global MAF ≥5% are shown separately by region, A: NPR; B: PAR1, C: PAR2; D: PAR3. For better visualization p-values < 1e-300 are plotted as 1e-300 (300 on -log10 scale). The red dashed line represents the line of equality. The corresponding Manhattan plots are in Figures S9 (across the whole X chromosome) and Figures S10, S11 and S12 for PAR1, PAR2 and PAR3, respectively.

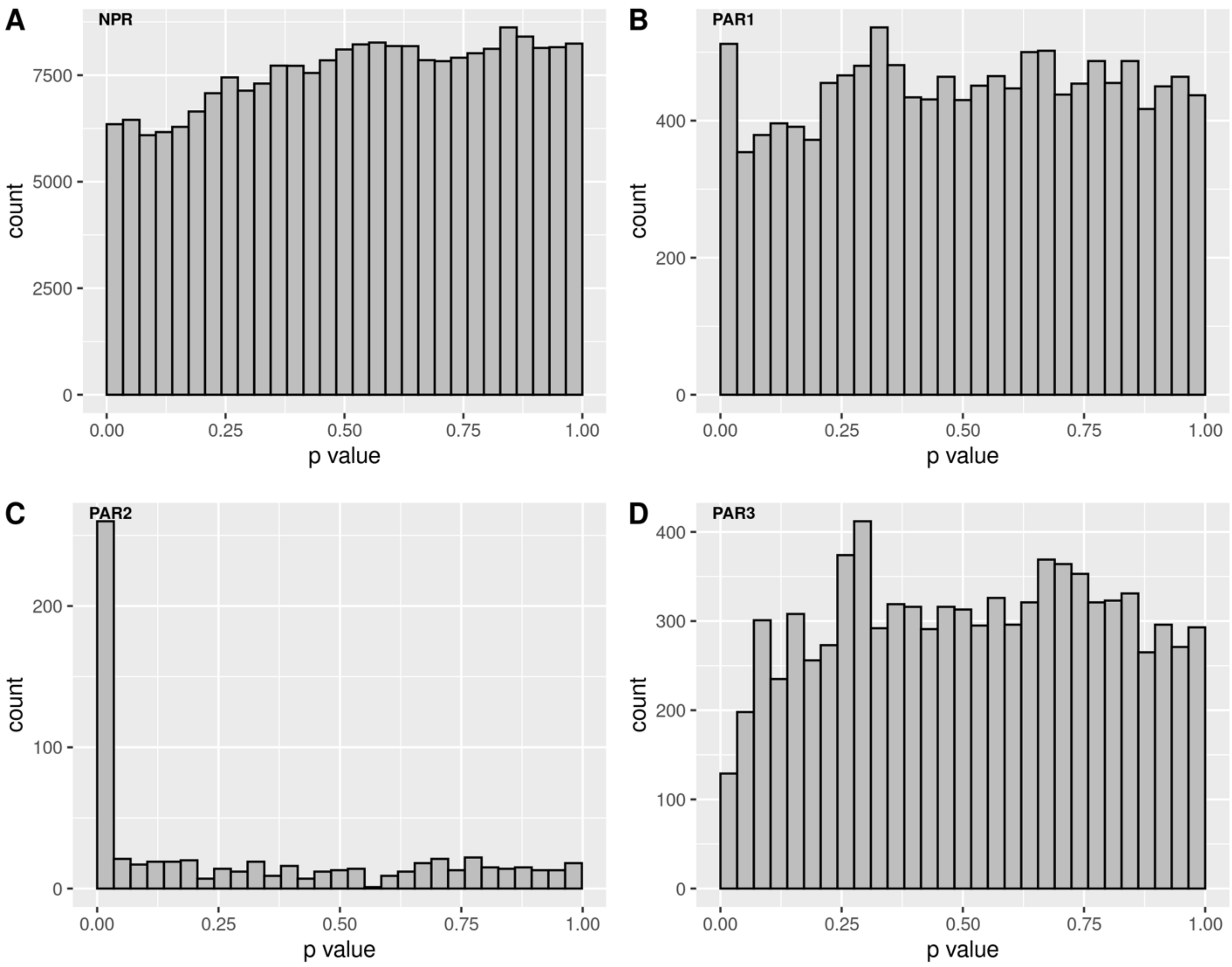


**Figure S14. Histograms of the sdMAF p-values of the X chromosome from the 1000 Genomes Project high coverage sequence data on GRCh38.** Results of bi-allelic SNPs with global MAF ≥5% are shown separately by region, A: NPR; B: PAR1, C: PAR2; D: PAR3. The corresponding Manhattan plots are in Figures S9 (across the whole X chromosome) and Figures S10, S11 and S12 for PAR1, PAR2 and PAR3, respectively.
