## Supplementary material for "Major sex differences in allele frequencies for X chromosome variants in the 1000 Genomes Project data": Note S1

<u>**Supplementary Note S1**</u>

**Bias in (naïve) population-pooled sample estimate of Hardy-Weinberg disequilibrium (HWD) across multiple populations.**

It is known that even if a bi-allelic SNP is in Hardy-Weinberg equilibrium (HWE) in each of the five super-populations or 26 populations, it may not be in HWE in the combined population, unless the MAFs are the same across all populations. Here we show that the bias factor for the naïve HWD estimate, when obtained from the population-pooled whole sample, is always greater or equal to zero.

Assume there are a total of $K$ populations, and let $\delta_k$ be the HWD in population $k, k = 1, ...K$, and $\hat{\delta}_k$ be the sample estimate from a sample size of $n_k$. For the bi-allelic SNP of interest, let $p_k(A)$ be the allele frequency of allele $A$ in population $k$ and $\hat{p}_k(A)$ the sample estimate, and let $p_k(AA)$ and $\hat{p}_k(AA)$ be the population genotype frequency of $AA$ and the corresponding sample estimate, respectively. Note that $\hat{p}_k(AA) = \hat{\delta}_k + \hat{p}_k(A)^2$, where $\hat{\delta}_k$, the population-stratified HWD estimate is defined as the difference between genotype $AA$ frequency estimate and squared of allele $A$ frequency estimate in population $k$.

The population-pooled sample estimates of allele and genotype frequencies are, respectively,

$$\hat{p}_w(A) = \frac{\sum_{k=1}^{K} \hat{p}_k(A) n_k}{\sum_{k=1}^{K} n_k}, \hat{p}_w(AA) = \frac{\sum_{k=1}^{K} \hat{p}_k(AA) n_k}{\sum_{k=1}^{K} n_k} = \frac{\sum_{k=1}^{K} \left(\hat{\delta}_k + \hat{p}_k(A)^2\right) n_k}{\sum_{k=1}^{K} n_k}.$$

The population-pooled sample estimate of HWD is then

$$\hat{\delta}_w = \frac{\sum_{k=1}^{K} \left(\hat{\delta}_k + \hat{p}_k(A)^2\right) n_k}{\sum_{k=1}^{K} n_k} - \left(\frac{\sum_{k=1}^{K} \hat{p}_k(A) n_k}{\sum_{k=1}^{K} n_k}\right)^2 = \frac{\sum_{k=1}^{K} \hat{\delta}_k n_k}{\sum_{k=1}^{K} n_k} + \frac{\left(\sum_{k=1}^{K} \hat{p}_k(A)^2 n_k\right)\left(\sum_{k=1}^{K} n_k\right) - \left(\sum_{k=1}^{K} \hat{p}_k(A) n_k\right)^2}{\left(\sum_{k=1}^{K} n_k\right)^2},$$

where the first term is a sample size-weighted linear combination of the population-stratified HWD estimates, $\hat{\delta}_k$, $k, k = 1, ...K$. If the SNP is in HWE in each of the $K$ individual populations, then $E[\hat{\delta}_k] = 0$ and $E\left[\frac{\sum_{k=1}^{K} \hat{\delta}_k n_k}{\sum_{k=1}^{K} n_k}\right] = \frac{\sum_{k=1}^{K} E[\hat{\delta}_k] n_k}{\sum_{k=1}^{K} n_k} = 0$.

Now we exam the numerator of the second term, $\left(\sum_{k=1}^{K} \hat{p}_k(A) n_k\right)^2$, in the expression for $\hat{\delta}_w$ above. Using the Cauchy–Schwarz inequality,

$$\left(\sum_{k=1}^{K} \hat{p}_k(A) n_k\right)^2 = \left(\sum_{k=1}^{K} \left(\hat{p}_k(A)\sqrt{n_k}\right)\left(\sqrt{n_k}\right)\right)^2$$

$$\leq \left(\sum_{k=1}^{K} \left(\hat{p}_k(A)\sqrt{n_k}\right)^2\right)\left(\sum_{k=1}^{K} \left(\sqrt{n_k}\right)^2\right)$$

$$= \left(\sum_{k=1}^{K} \hat{p}_k(A)^2 n_k\right)\left(\sum_{k=1}^{K} n_k\right).$$

Thus, the second term for $\hat{\delta}_w$ is always greater or equal to zero. This explains why we performed the HWE test separately for each of the five super-populations. Valid HWD testing across multiple populations is possible (1) and (2), but the population-pooled approach is not suitable for the purpose of our HWD analysis, which examines whether HWD is present in a specific super-population.

For the test of sex differences in minor allele frequency (sdMAF), however, the population-pooled whole-sample approach is more powerful and also valid, because the sdMAF test detects the difference in MAF between males and females, not the difference in MAF between populations. However, the bias in the naïve population-pooled sample estimate of HWD leads to a bigger variance, which in turn leads to a conservative sdMAF test.
