## Supplementary material for "Major sex differences in allele frequencies for X chromosome variants in the 1000 Genomes Project data": Note S2

### Supplementary Note 2

**Comparison of genotypes for selected SNPs between phase 3 and high coverage whole genome sequence data from the 1000 Genomes Project.**

In total, 10, 50, 20, and 50 SNPs, respectively from NPR, PAR1, PAR2, and PAR3, with the smallest sdMAF p-values in the phase 3 data were first selected.

Among these SNPs, 10, 4, 9, and 9 SNPs, respectively from NPR, PAR1, PAR2, and PAR3, were also bi-allelic in the high coverage data and had no missingness in both sets of data.

Each page represents the results for one SNP, and SNPs are ordered by the GRCh37 positions.

Within each page, the position of the SNP in phase 3 (build GRCh37) and high coverage (GRCh38) are first provided. Next is the female - male sdMAF difference and the sdMAF p-value. The REF and ALT alleles are also provided for each build. Finally, the counts of the agreement of the genotype calls between the phase 3 and the high coverage data are provided, separately by sex.

Region: PAR1
rs number (GRCh37 phase3): .
SNP position (GRCh37) = 2695860 ; Female-Male Sex Difference in MAF (GRCh37 phase3) = -0.3727 ; sdMAF P-value = 8.20918418869952e-268
SNP position (GRCh38) = 2777819 ; Female-Male Sex Difference in MAF (High Coverage) = -0.3739 ; sdMAF P-value = 7.863953925689e-273

Table 1: REF/ALT

| | REF | ALT |
|---|---|---|
| Phase3 (GRCh37) | A | G |
| High Coverage | A | G |

Table 2: male

| | aa | Aa | AA | Total_P3 |
|---|---|---|---|---|
| aa | 99 | 3 | 0 | 102 |
| Aa | 0 | 917 | 0 | 917 |
| AA | 0 | 1 | 213 | 214 |
| Total_HC | 99 | 921 | 213 | 1233 |

Table 3: female

| | aa | Aa | AA | Total_P3 |
|---|---|---|---|---|
| aa | 892 | 1 | 0 | 893 |
| Aa | 0 | 317 | 0 | 317 |
| AA | 0 | 2 | 59 | 61 |
| Total_HC | 892 | 320 | 59 | 1271 |

Note1. Row (Phase 3 GRCh37); Column (High Coverage GRCh38)
Note2. ‘a’: REF; ‘A’: ALT

Region: PAR1
rs number (GRCh37 phase3): .
SNP position (GRCh37) = 2696560 ; Female-Male Sex Difference in MAF (GRCh37 phase3) = -0.376 ; sdMAF P-value = 5.54370357180522e-289
SNP position (GRCh38) = 2778519 ; Female-Male Sex Difference in MAF (High Coverage) = -0.3748 ; sdMAF P-value = 2.65832322082409e-287

Table 4: REF/ALT

| | REF | ALT |
|---|---|---|
| Phase3 (GRCh37) | G | A |
| High Coverage | G | A |

Table 5: male

| | aa | Aa | AA | Total_P3 |
|---|---|---|---|---|
| aa | 98 | 1 | 0 | 99 |
| Aa | 0 | 940 | 0 | 940 |
| AA | 0 | 2 | 192 | 194 |
| Total_HC | 98 | 943 | 192 | 1233 |

Table 6: female

| | aa | Aa | AA | Total_P3 |
|---|---|---|---|---|
| aa | 912 | 1 | 0 | 913 |
| Aa | 0 | 302 | 1 | 303 |
| AA | 0 | 0 | 55 | 55 |
| Total_HC | 912 | 303 | 56 | 1271 |

Note1. Row (Phase 3 GRCh37); Column (High Coverage GRCh38)
Note2. ‘a’: REF; ‘A’: ALT

Region: PAR1
rs number (GRCh37 phase3): .
SNP position (GRCh37) = 2696892 ; Female-Male Sex Difference in MAF (GRCh37 phase3) = -0.4001 ; sdMAF P-value < 1e-300
SNP position (GRCh38) = 2778851 ; Female-Male Sex Difference in MAF (High Coverage) = -0.40047 ; sdMAF P-value < 1e-300

Table 7: REF/ALT

| | REF | ALT |
|---|---|---|
| Phase3 (GRCh37) | G | A |
| High Coverage | G | A |

Table 8: male

| | aa | Aa | AA | Total_P3 |
|---|---|---|---|---|
| aa | 103 | 2 | 0 | 105 |
| Aa | 1 | 1019 | 0 | 1020 |
| AA | 0 | 4 | 104 | 108 |
| Total_HC | 104 | 1025 | 104 | 1233 |

Table 9: female

| | aa | Aa | AA | Total_P3 |
|---|---|---|---|---|
| aa | 1041 | 0 | 0 | 1041 |
| Aa | 2 | 201 | 0 | 203 |
| AA | 0 | 2 | 25 | 27 |
| Total_HC | 1043 | 203 | 25 | 1271 |

Note1. Row (Phase 3 GRCh37); Column (High Coverage GRCh38)
Note2. ‘a’: REF; ‘A’: ALT

Region: PAR1
rs number (GRCh37 phase3): .
SNP position (GRCh37) = 2696893 ; Female-Male Sex Difference in MAF (GRCh37 phase3) = -0.4001 ; sdMAF P-value < 1e-300
SNP position (GRCh38) = 2778852 ; Female-Male Sex Difference in MAF (High Coverage) = -0.40047 ; sdMAF P-value < 1e-300

Table 10: REF/ALT

| | REF | ALT |
|---|---|---|
| Phase3 (GRCh37) | C | T |
| High Coverage | C | T |

Table 11: male

| | aa | Aa | AA | Total_P3 |
|---|---|---|---|---|
| aa | 103 | 2 | 0 | 105 |
| Aa | 1 | 1019 | 0 | 1020 |
| AA | 0 | 4 | 104 | 108 |
| Total_HC | 104 | 1025 | 104 | 1233 |

Table 12: female

| | aa | Aa | AA | Total_P3 |
|---|---|---|---|---|
| aa | 1041 | 0 | 0 | 1041 |
| Aa | 2 | 201 | 0 | 203 |
| AA | 0 | 2 | 25 | 27 |
| Total_HC | 1043 | 203 | 25 | 1271 |

Note1. Row (Phase 3 GRCh37); Column (High Coverage GRCh38)
Note2. ‘a’: REF; ‘A’: ALT

Region: PAR1
rs number (GRCh37 phase3): .
SNP position (GRCh37) = 2697154 ; Female-Male Sex Difference in MAF (GRCh37 phase3) = -0.41948 ; sdMAF P-value < 1e-300
SNP position (GRCh38) = 2779113 ; Female-Male Sex Difference in MAF (High Coverage) = -0.4183 ; sdMAF P-value < 1e-300

Table 13: REF/ALT

| | REF | ALT |
|---|---|---|
| Phase3 (GRCh37) | A | C |
| High Coverage | A | C |

Table 14: male

| | aa | Aa | AA | Total_P3 |
|---|---|---|---|---|
| aa | 106 | 2 | 0 | 108 |
| Aa | 1 | 1040 | 2 | 1043 |
| AA | 0 | 3 | 79 | 82 |
| Total_HC | 107 | 1045 | 81 | 1233 |

Table 15: female

| | aa | Aa | AA | Total_P3 |
|---|---|---|---|---|
| aa | 1097 | 3 | 0 | 1100 |
| Aa | 1 | 162 | 1 | 164 |
| AA | 0 | 0 | 7 | 7 |
| Total_HC | 1098 | 165 | 8 | 1271 |

Note1. Row (Phase 3 GRCh37); Column (High Coverage GRCh38)
Note2. ‘a’: REF; ‘A’: ALT

Region: PAR1
rs number (GRCh37 phase3): .
SNP position (GRCh37) = 2697599 ; Female-Male Sex Difference in MAF (GRCh37 phase3) = -0.461312 ; sdMAF P-value < 1e-300
SNP position (GRCh38) = 2779558 ; Female-Male Sex Difference in MAF (High Coverage) = -0.458793 ; sdMAF P-value < 1e-300

Table 16: REF/ALT

| | REF | ALT |
|---|---|---|
| Phase3 (GRCh37) | C | A |
| High Coverage | C | A |

Table 17: male

| | aa | Aa | AA | Total_P3 |
|---|---|---|---|---|
| aa | 88 | 2 | 0 | 90 |
| Aa | 7 | 1125 | 0 | 1132 |
| AA | 0 | 4 | 7 | 11 |
| Total_HC | 95 | 1131 | 7 | 1233 |

Table 18: female

| | aa | Aa | AA | Total_P3 |
|---|---|---|---|---|
| aa | 1253 | 1 | 0 | 1254 |
| Aa | 4 | 13 | 0 | 17 |
| AA | 0 | 0 | 0 | 0 |
| Total_HC | 1257 | 14 | 0 | 1271 |

Note1. Row (Phase 3 GRCh37); Column (High Coverage GRCh38)
Note2. ‘a’: REF; ‘A’: ALT

Region: PAR1
rs number (GRCh37 phase3): .
SNP position (GRCh37) = 2697845 ; Female-Male Sex Difference in MAF (GRCh37 phase3) = -0.42742 ; sdMAF P-value < 1e-300
SNP position (GRCh38) = 2779804 ; Female-Male Sex Difference in MAF (High Coverage) = -0.42582 ; sdMAF P-value < 1e-300

Table 19: REF/ALT

| | REF | ALT |
|---|---|---|
| Phase3 (GRCh37) | C | T |
| High Coverage | C | T |

Table 20: male

| | aa | Aa | AA | Total_P3 |
|---|---|---|---|---|
| aa | 78 | 2 | 0 | 80 |
| Aa | 1 | 1057 | 1 | 1059 |
| AA | 0 | 5 | 89 | 94 |
| Total_HC | 79 | 1064 | 90 | 1233 |

Table 21: female

| | aa | Aa | AA | Total_P3 |
|---|---|---|---|---|
| aa | 1089 | 4 | 0 | 1093 |
| Aa | 2 | 155 | 0 | 157 |
| AA | 0 | 1 | 20 | 21 |
| Total_HC | 1091 | 160 | 20 | 1271 |

Note1. Row (Phase 3 GRCh37); Column (High Coverage GRCh38)
Note2. ‘a’: REF; ‘A’: ALT

Region: PAR1
rs number (GRCh37 phase3): .
SNP position (GRCh37) = 2697868 ; Female-Male Sex Difference in MAF (GRCh37 phase3) = -0.43172 ; sdMAF P-value < 1e-300
SNP position (GRCh38) = 2779827 ; Female-Male Sex Difference in MAF (High Coverage) = -0.43133 ; sdMAF P-value < 1e-300

Table 22: REF/ALT

| | REF | ALT |
|---|---|---|
| Phase3 (GRCh37) | G | A |
| High Coverage | G | A |

Table 23: male

| | aa | Aa | AA | Total_P3 |
|---|---|---|---|---|
| aa | 78 | 2 | 0 | 80 |
| Aa | 0 | 1059 | 3 | 1062 |
| AA | 0 | 5 | 86 | 91 |
| Total_HC | 78 | 1066 | 89 | 1233 |

Table 24: female

| | aa | Aa | AA | Total_P3 |
|---|---|---|---|---|
| aa | 1100 | 4 | 0 | 1104 |
| Aa | 2 | 147 | 0 | 149 |
| AA | 0 | 1 | 17 | 18 |
| Total_HC | 1102 | 152 | 17 | 1271 |

Note1. Row (Phase 3 GRCh37); Column (High Coverage GRCh38)
Note2. ‘a’: REF; ‘A’: ALT

Region: PAR1
rs number (GRCh37 phase3): .
SNP position (GRCh37) = 2698923 ; Female-Male Sex Difference in MAF (GRCh37 phase3) = -0.46516 ; sdMAF P-value < 1e-300
SNP position (GRCh38) = 2780882 ; Female-Male Sex Difference in MAF (High Coverage) = -0.46225 ; sdMAF P-value < 1e-300

Table 25: REF/ALT

| | REF | ALT |
|---|---|---|
| Phase3 (GRCh37) | G | A |
| High Coverage | G | A |

Table 26: male

| | aa | Aa | AA | Total_P3 |
|---|---|---|---|---|
| aa | 38 | 6 | 0 | 44 |
| Aa | 6 | 1128 | 0 | 1134 |
| AA | 0 | 8 | 47 | 55 |
| Total_HC | 44 | 1142 | 47 | 1233 |

Table 27: female

| | aa | Aa | AA | Total_P3 |
|---|---|---|---|---|
| aa | 1178 | 0 | 0 | 1178 |
| Aa | 0 | 85 | 1 | 86 |
| AA | 0 | 2 | 5 | 7 |
| Total_HC | 1178 | 87 | 6 | 1271 |

Note1. Row (Phase 3 GRCh37); Column (High Coverage GRCh38)
Note2. ‘a’: REF; ‘A’: ALT

Region: PAR1
rs number (GRCh37 phase3): .
SNP position (GRCh37) = 2698954 ; Female-Male Sex Difference in MAF (GRCh37 phase3) = -0.427 ; sdMAF P-value < 1e-300
SNP position (GRCh38) = 2780913 ; Female-Male Sex Difference in MAF (High Coverage) = -0.4245 ; sdMAF P-value < 1e-300

Table 28: REF/ALT

| | REF | ALT |
|---|---|---|
| Phase3 (GRCh37) | G | A |
| High Coverage | G | A |

Table 29: male

| | aa | Aa | AA | Total_P3 |
|---|---|---|---|---|
| aa | 37 | 5 | 0 | 42 |
| Aa | 1 | 1011 | 1 | 1013 |
| AA | 0 | 10 | 168 | 178 |
| Total_HC | 38 | 1026 | 169 | 1233 |

Table 30: female

| | aa | Aa | AA | Total_P3 |
|---|---|---|---|---|
| aa | 975 | 4 | 0 | 979 |
| Aa | 3 | 254 | 1 | 258 |
| AA | 0 | 1 | 33 | 34 |
| Total_HC | 978 | 259 | 34 | 1271 |

Note1. Row (Phase 3 GRCh37); Column (High Coverage GRCh38)
Note2. ‘a’: REF; ‘A’: ALT

Region: NPR
rs number (GRCh37 phase3): rs1996225
SNP position (GRCh37) = 15711209 ; Female-Male Sex Difference in MAF (GRCh37 phase3) = 0.2344 ; sdMAF P-value = 1.55380206985061e-60
SNP position (GRCh38) = 15693086 ; Female-Male Sex Difference in MAF (High Coverage) = 0.0352 ; sdMAF P-value = 0.0251232744816044

Table 31: REF/ALT

| | REF | ALT |
|---|---|---|
| Phase3 (GRCh37) | C | T |
| High Coverage | C | T |

Table 32: male

| | a | A | Total_P3 |
|---|---|---|---|
| a | 315 | 11 | 326 |
| A | 0 | 907 | 907 |
| Total_HC | 315 | 918 | 1233 |

Table 33: female

| | aa | Aa | AA | Total_P3 |
|---|---|---|---|---|
| aa | 147 | 1 | 0 | 148 |
| Aa | 0 | 444 | 528 | 972 |
| AA | 0 | 0 | 151 | 151 |
| Total_HC | 147 | 445 | 679 | 1271 |

Note1. Row (Phase 3 GRCh37); Column (High Coverage GRCh38)
Note2. ‘a’: REF; ‘A’: ALT

Region: NPR
rs number (GRCh37 phase3): rs372984882
SNP position (GRCh37) = 88383111 ; Female-Male Sex Difference in MAF (GRCh37 phase3) = 0.28941 ; sdMAF P-value = 3.9737450506146e-213
SNP position (GRCh38) = 89128111 ; Female-Male Sex Difference in MAF (High Coverage) = 0.00236 ; sdMAF P-value = 0.0140770215674526

Table 34: REF/ALT

| | REF | ALT |
|---|---|---|
| Phase3 (GRCh37) | A | T |
| High Coverage | A | T |

Table 35: male

| | a | A | Total_P3 |
|---|---|---|---|
| a | 1161 | 0 | 1161 |
| A | 72 | 0 | 72 |
| Total_HC | 1233 | 0 | 1233 |

Table 36: female

| | aa | Aa | AA | Total_P3 |
|---|---|---|---|---|
| aa | 384 | 3 | 0 | 387 |
| Aa | 881 | 3 | 0 | 884 |
| AA | 0 | 0 | 0 | 0 |
| Total_HC | 1265 | 6 | 0 | 1271 |

Note1. Row (Phase 3 GRCh37); Column (High Coverage GRCh38)
Note2. ‘a’: REF; ‘A’: ALT

Region: NPR
rs number (GRCh37 phase3): rs369028615
SNP position (GRCh37) = 88383115 ; Female-Male Sex Difference in MAF (GRCh37 phase3) = 0.28941 ; sdMAF P-value = 3.9737450506146e-213
SNP position (GRCh38) = 89128115 ; Female-Male Sex Difference in MAF (High Coverage) = 0.00236 ; sdMAF P-value = 0.0140770215674526

Table 37: REF/ALT

| | REF | ALT |
|---|---|---|
| Phase3 (GRCh37) | C | T |
| High Coverage | C | T |

Table 38: male

| | a | A | Total_P3 |
|---|---|---|---|
| a | 1161 | 0 | 1161 |
| A | 72 | 0 | 72 |
| Total_HC | 1233 | 0 | 1233 |

Table 39: female

| | aa | Aa | AA | Total_P3 |
|---|---|---|---|---|
| aa | 384 | 3 | 0 | 387 |
| Aa | 881 | 3 | 0 | 884 |
| AA | 0 | 0 | 0 | 0 |
| Total_HC | 1265 | 6 | 0 | 1271 |

Note1. Row (Phase 3 GRCh37); Column (High Coverage GRCh38)
Note2. ‘a’: REF; ‘A’: ALT

Region: PAR3
rs number (GRCh37 phase3): .
SNP position (GRCh37) = 88458816 ; Female-Male Sex Difference in MAF (GRCh37 phase3) = 0.16151 ; sdMAF P-value = 2.25662814279251e-56
SNP position (GRCh38) = 89203817 ; Female-Male Sex Difference in MAF (High Coverage) = 0.04013 ; sdMAF P-value = 6.22346860302191e-26

Table 40: REF/ALT

| | REF | ALT |
|---|---|---|
| Phase3 (GRCh37) | G | A |
| High Coverage | G | A |

Table 41: male

| | a | A | Total_P3 |
|---|---|---|---|
| a | 1143 | 0 | 1143 |
| A | 90 | 0 | 90 |
| Total_HC | 1233 | 0 | 1233 |

Table 42: female

| | aa | Aa | AA | Total_P3 |
|---|---|---|---|---|
| aa | 649 | 27 | 0 | 676 |
| Aa | 519 | 75 | 0 | 594 |
| AA | 1 | 0 | 0 | 1 |
| Total_HC | 1169 | 102 | 0 | 1271 |

Note1. Row (Phase 3 GRCh37); Column (High Coverage GRCh38)
Note2. ‘a’: REF; ‘A’: ALT

Region: PAR3
rs number (GRCh37 phase3): .
SNP position (GRCh37) = 88458918 ; Female-Male Sex Difference in MAF (GRCh37 phase3) = 0.1504 ; sdMAF P-value = 3.10020205638595e-48
SNP position (GRCh38) = 89203919 ; Female-Male Sex Difference in MAF (High Coverage) = 0.04917 ; sdMAF P-value = 5.29484546944359e-32

Table 43: REF/ALT

| | REF | ALT |
|---|---|---|
| Phase3 (GRCh37) | A | G |
| High Coverage | A | G |

Table 44: male

| | a | A | Total_P3 |
|---|---|---|---|
| a | 1142 | 0 | 1142 |
| A | 91 | 0 | 91 |
| Total_HC | 1233 | 0 | 1233 |

Table 45: female

| | aa | Aa | AA | Total_P3 |
|---|---|---|---|---|
| aa | 651 | 57 | 0 | 708 |
| Aa | 490 | 66 | 0 | 556 |
| AA | 5 | 2 | 0 | 7 |
| Total_HC | 1146 | 125 | 0 | 1271 |

Note1. Row (Phase 3 GRCh37); Column (High Coverage GRCh38)
Note2. ‘a’: REF; ‘A’: ALT

Region: PAR3
rs number (GRCh37 phase3): rs370154167
SNP position (GRCh37) = 88460271 ; Female-Male Sex Difference in MAF (GRCh37 phase3) = 0.2629 ; sdMAF P-value = 9.87188301068426e-132
SNP position (GRCh38) = 89205272 ; Female-Male Sex Difference in MAF (High Coverage) = 0.002754 ; sdMAF P-value = 0.00797639777080226

Table 46: REF/ALT

| | REF | ALT |
|---|---|---|
| Phase3 (GRCh37) | G | A |
| High Coverage | G | A |

Table 47: male

| | a | A | Total_P3 |
|---|---|---|---|
| a | 1108 | 0 | 1108 |
| A | 125 | 0 | 125 |
| Total_HC | 1233 | 0 | 1233 |

Table 48: female

| | aa | Aa | AA | Total_P3 |
|---|---|---|---|---|
| aa | 354 | 1 | 0 | 355 |
| Aa | 900 | 6 | 0 | 906 |
| AA | 10 | 0 | 0 | 10 |
| Total_HC | 1264 | 7 | 0 | 1271 |

Note1. Row (Phase 3 GRCh37); Column (High Coverage GRCh38)
Note2. ‘a’: REF; ‘A’: ALT

Region: PAR3
rs number (GRCh37 phase3): .
SNP position (GRCh37) = 88460457 ; Female-Male Sex Difference in MAF (GRCh37 phase3) = 0.125 ; sdMAF P-value = 3.85096879960068e-45
SNP position (GRCh38) = 89205458 ; Female-Male Sex Difference in MAF (High Coverage) = -2.42e-05 ; sdMAF P-value = 0.980320072234104

Table 49: REF/ALT

| | REF | ALT |
|---|---|---|
| Phase3 (GRCh37) | G | A |
| High Coverage | G | C |

Table 50: male

| | a | A | Total_P3 |
|---|---|---|---|
| a | 1178 | 1 | 1179 |
| A | 54 | 0 | 54 |
| Total_HC | 1232 | 1 | 1233 |

Table 51: female

| | aa | Aa | AA | Total_P3 |
|---|---|---|---|---|
| aa | 842 | 2 | 0 | 844 |
| Aa | 425 | 0 | 0 | 425 |
| AA | 2 | 0 | 0 | 2 |
| Total_HC | 1269 | 2 | 0 | 1271 |

Note1. Row (Phase 3 GRCh37); Column (High Coverage GRCh38)
Note2. ‘a’: REF; ‘A’: ALT

Region: PAR3
rs number (GRCh37 phase3): .
SNP position (GRCh37) = 88460883 ; Female-Male Sex Difference in MAF (GRCh37 phase3) = 0.0985 ; sdMAF P-value = 3.54405130280993e-13
SNP position (GRCh38) = 89205884 ; Female-Male Sex Difference in MAF (High Coverage) = 0.015911 ; sdMAF P-value = 3.74164715957335e-05

Table 52: REF/ALT

| | REF | ALT |
|---|---|---|
| Phase3 (GRCh37) | G | A |
| High Coverage | G | A |

Table 53: male

| | a | A | Total_P3 |
|---|---|---|---|
| a | 978 | 5 | 983 |
| A | 246 | 4 | 250 |
| Total_HC | 1224 | 9 | 1233 |

Table 54: female

| | aa | Aa | AA | Total_P3 |
|---|---|---|---|---|
| aa | 504 | 19 | 0 | 523 |
| Aa | 692 | 37 | 1 | 730 |
| AA | 17 | 1 | 0 | 18 |
| Total_HC | 1213 | 57 | 1 | 1271 |

Note1. Row (Phase 3 GRCh37); Column (High Coverage GRCh38)
Note2. ‘a’: REF; ‘A’: ALT

Region: PAR3
rs number (GRCh37 phase3): rs199820323
SNP position (GRCh37) = 89192798 ; Female-Male Sex Difference in MAF (GRCh37 phase3) = 0.10024 ; sdMAF P-value = 4.78641113314677e-32
SNP position (GRCh38) = 89937799 ; Female-Male Sex Difference in MAF (High Coverage) = -0.00602 ; sdMAF P-value = 0.157584761057514

Table 55: REF/ALT

| | REF | ALT |
|---|---|---|
| Phase3 (GRCh37) | C | A |
| High Coverage | C | A |

Table 56: male

| | a | A | Total_P3 |
|---|---|---|---|
| a | 1182 | 0 | 1182 |
| A | 30 | 21 | 51 |
| Total_HC | 1212 | 21 | 1233 |

Table 57: female

| | aa | Aa | AA | Total_P3 |
|---|---|---|---|---|
| aa | 911 | 1 | 0 | 912 |
| Aa | 333 | 25 | 0 | 358 |
| AA | 0 | 0 | 1 | 1 |
| Total_HC | 1244 | 26 | 1 | 1271 |

Note1. Row (Phase 3 GRCh37); Column (High Coverage GRCh38)
Note2. ‘a’: REF; ‘A’: ALT

Region: PAR3
rs number (GRCh37 phase3): rs113948071
SNP position (GRCh37) = 89803933 ; Female-Male Sex Difference in MAF (GRCh37 phase3) = 0.12154 ; sdMAF P-value = 2.45504806968182e-44
SNP position (GRCh38) = 90548934 ; Female-Male Sex Difference in MAF (High Coverage) = -0.000811 ; sdMAF P-value = 0.31711418273145

Table 58: REF/ALT

| | REF | ALT |
|---|---|---|
| Phase3 (GRCh37) | A | C |
| High Coverage | A | C |

Table 59: male

| | a | A | Total_P3 |
|---|---|---|---|
| a | 1182 | 0 | 1182 |
| A | 50 | 1 | 51 |
| Total_HC | 1232 | 1 | 1233 |

Table 60: female

| | aa | Aa | AA | Total_P3 |
|---|---|---|---|---|
| aa | 858 | 0 | 0 | 858 |
| Aa | 412 | 0 | 0 | 412 |
| AA | 1 | 0 | 0 | 1 |
| Total_HC | 1271 | 0 | 0 | 1271 |

Note1. Row (Phase 3 GRCh37); Column (High Coverage GRCh38)
Note2. ‘a’: REF; ‘A’: ALT

Region: PAR3
rs number (GRCh37 phase3): rs75914390
SNP position (GRCh37) = 90420559 ; Female-Male Sex Difference in MAF (GRCh37 phase3) = 0.13493 ; sdMAF P-value = 1.83772278515363e-34
SNP position (GRCh38) = 91165560 ; Female-Male Sex Difference in MAF (High Coverage) = 0.00849 ; sdMAF P-value = 0.308025448569295

Table 61: REF/ALT

| | REF | ALT |
|---|---|---|
| Phase3 (GRCh37) | C | G |
| High Coverage | C | G |

Table 62: male

| | a | A | Total_P3 |
|---|---|---|---|
| a | 1118 | 0 | 1118 |
| A | 43 | 72 | 115 |
| Total_HC | 1161 | 72 | 1233 |

Table 63: female

| | aa | Aa | AA | Total_P3 |
|---|---|---|---|---|
| aa | 704 | 0 | 0 | 704 |
| Aa | 402 | 152 | 0 | 554 |
| AA | 1 | 6 | 6 | 13 |
| Total_HC | 1107 | 158 | 6 | 1271 |

Note1. Row (Phase 3 GRCh37); Column (High Coverage GRCh38)
Note2. ‘a’: REF; ‘A’: ALT

Region: PAR3
rs number (GRCh37 phase3): rs56118643
SNP position (GRCh37) = 90471772 ; Female-Male Sex Difference in MAF (GRCh37 phase3) = -0.0863 ; sdMAF P-value = 7.46065572919066e-08
SNP position (GRCh38) = 91216773 ; Female-Male Sex Difference in MAF (High Coverage) = -0.0031 ; sdMAF P-value = 0.853222915843523

Table 64: REF/ALT

| | REF | ALT |
|---|---|---|
| Phase3 (GRCh37) | T | A |
| High Coverage | T | A |

Table 65: male

| | a | A | Total_P3 |
|---|---|---|---|
| a | 812 | 55 | 867 |
| A | 1 | 365 | 366 |
| Total_HC | 813 | 420 | 1233 |

Table 66: female

| | aa | Aa | AA | Total_P3 |
|---|---|---|---|---|
| aa | 584 | 275 | 8 | 867 |
| Aa | 10 | 221 | 42 | 273 |
| AA | 0 | 0 | 131 | 131 |
| Total_HC | 594 | 496 | 181 | 1271 |

Note1. Row (Phase 3 GRCh37); Column (High Coverage GRCh38)
Note2. ‘a’: REF; ‘A’: ALT

Region: NPR
rs number (GRCh37 phase3): rs6637609
SNP position (GRCh37) = 128638559 ; Female-Male Sex Difference in MAF (GRCh37 phase3) = 0.2211 ; sdMAF P-value = 6.54681396294046e-63
SNP position (GRCh38) = 129504582 ; Female-Male Sex Difference in MAF (High Coverage) = 0.021 ; sdMAF P-value = 0.0770279808229025

Table 67: REF/ALT

| | REF | ALT |
|---|---|---|
| Phase3 (GRCh37) | C | A |
| High Coverage | C | A |

Table 68: male

| | a | A | Total_P3 |
|---|---|---|---|
| a | 920 | 3 | 923 |
| A | 157 | 153 | 310 |
| Total_HC | 1077 | 156 | 1233 |

Table 69: female

| | aa | Aa | AA | Total_P3 |
|---|---|---|---|---|
| aa | 102 | 5 | 0 | 107 |
| Aa | 826 | 300 | 1 | 1127 |
| AA | 2 | 2 | 33 | 37 |
| Total_HC | 930 | 307 | 34 | 1271 |

Note1. Row (Phase 3 GRCh37); Column (High Coverage GRCh38)
Note2. ‘a’: REF; ‘A’: ALT

Region: PAR2
rs number (GRCh37 phase3): .
SNP position (GRCh37) = 154934428 ; Female-Male Sex Difference in MAF (GRCh37 phase3) = 0.3434 ; sdMAF P-value = 1.22828740905784e-169
SNP position (GRCh38) = 155704767 ; Female-Male Sex Difference in MAF (High Coverage) = 0.3442 ; sdMAF P-value = 5.16020258391841e-171

Table 70: REF/ALT

| | REF | ALT |
|---|---|---|
| Phase3 (GRCh37) | T | G |
| High Coverage | T | G |

Table 71: male

| | aa | Aa | AA | Total_P3 |
|---|---|---|---|---|
| aa | 1 | 1 | 0 | 2 |
| Aa | 0 | 745 | 0 | 745 |
| AA | 0 | 0 | 486 | 486 |
| Total_HC | 1 | 746 | 486 | 1233 |

Table 72: female

| | aa | Aa | AA | Total_P3 |
|---|---|---|---|---|
| aa | 578 | 0 | 0 | 578 |
| Aa | 0 | 489 | 0 | 489 |
| AA | 0 | 1 | 203 | 204 |
| Total_HC | 578 | 490 | 203 | 1271 |

Note1. Row (Phase 3 GRCh37); Column (High Coverage GRCh38)
Note2. ‘a’: REF; ‘A’: ALT

Region: PAR2
rs number (GRCh37 phase3): .
SNP position (GRCh37) = 154934986 ; Female-Male Sex Difference in MAF (GRCh37 phase3) = 0.3433 ; sdMAF P-value = 4.76760207367504e-169
SNP position (GRCh38) = 155705325 ; Female-Male Sex Difference in MAF (High Coverage) = 0.3458 ; sdMAF P-value = 5.65681734446808e-173

Table 73: REF/ALT

| | REF | ALT |
|---|---|---|
| Phase3 (GRCh37) | G | A |
| High Coverage | G | A |

Table 74: male

| | aa | Aa | AA | Total_P3 |
|---|---|---|---|---|
| aa | 1 | 5 | 0 | 6 |
| Aa | 0 | 741 | 0 | 741 |
| AA | 0 | 0 | 486 | 486 |
| Total_HC | 1 | 746 | 486 | 1233 |

Table 75: female

| | aa | Aa | AA | Total_P3 |
|---|---|---|---|---|
| aa | 578 | 1 | 0 | 579 |
| Aa | 2 | 489 | 0 | 491 |
| AA | 0 | 0 | 201 | 201 |
| Total_HC | 580 | 490 | 201 | 1271 |

Note1. Row (Phase 3 GRCh37); Column (High Coverage GRCh38)
Note2. ‘a’: REF; ‘A’: ALT

Region: PAR2
rs number (GRCh37 phase3): .
SNP position (GRCh37) = 154938547 ; Female-Male Sex Difference in MAF (GRCh37 phase3) = 0.343 ; sdMAF P-value = 2.93224328230306e-169
SNP position (GRCh38) = 155708886 ; Female-Male Sex Difference in MAF (High Coverage) = 0.3434 ; sdMAF P-value = 2.95168132842575e-169

Table 76: REF/ALT

| | REF | ALT |
|---|---|---|
| Phase3 (GRCh37) | T | C |
| High Coverage | T | C |

Table 77: male

| | aa | Aa | AA | Total_P3 |
|---|---|---|---|---|
| aa | 2 | 1 | 0 | 3 |
| Aa | 0 | 744 | 0 | 744 |
| AA | 0 | 1 | 485 | 486 |
| Total_HC | 2 | 746 | 485 | 1233 |

Table 78: female

| | aa | Aa | AA | Total_P3 |
|---|---|---|---|---|
| aa | 577 | 0 | 0 | 577 |
| Aa | 3 | 486 | 2 | 491 |
| AA | 0 | 0 | 203 | 203 |
| Total_HC | 580 | 486 | 205 | 1271 |

Note1. Row (Phase 3 GRCh37); Column (High Coverage GRCh38)
Note2. ‘a’: REF; ‘A’: ALT

Region: PAR2
rs number (GRCh37 phase3): .
SNP position (GRCh37) = 154950587 ; Female-Male Sex Difference in MAF (GRCh37 phase3) = -0.4092 ; sdMAF P-value < 1e-300
SNP position (GRCh38) = 155720925 ; Female-Male Sex Difference in MAF (High Coverage) = -0.41 ; sdMAF P-value < 1e-300

Table 79: REF/ALT

| | REF | ALT |
|---|---|---|
| Phase3 (GRCh37) | T | C |
| High Coverage | T | C |

Table 80: male

| | aa | Aa | AA | Total_P3 |
|---|---|---|---|---|
| aa | 21 | 1 | 0 | 22 |
| Aa | 0 | 953 | 1 | 954 |
| AA | 0 | 0 | 257 | 257 |
| Total_HC | 21 | 954 | 258 | 1233 |

Table 81: female

| | aa | Aa | AA | Total_P3 |
|---|---|---|---|---|
| aa | 859 | 0 | 0 | 859 |
| Aa | 0 | 349 | 2 | 351 |
| AA | 0 | 2 | 59 | 61 |
| Total_HC | 859 | 351 | 61 | 1271 |

Note1. Row (Phase 3 GRCh37); Column (High Coverage GRCh38)
Note2. ‘a’: REF; ‘A’: ALT

Region: PAR2
rs number (GRCh37 phase3): .
SNP position (GRCh37) = 154954348 ; Female-Male Sex Difference in MAF (GRCh37 phase3) = -0.408 ; sdMAF P-value < 1e-300
SNP position (GRCh38) = 155724686 ; Female-Male Sex Difference in MAF (High Coverage) = -0.4084 ; sdMAF P-value < 1e-300

Table 82: REF/ALT

| | REF | ALT |
|---|---|---|
| Phase3 (GRCh37) | C | T |
| High Coverage | C | T |

Table 83: male

| | aa | Aa | AA | Total_P3 |
|---|---|---|---|---|
| aa | 23 | 1 | 0 | 24 |
| Aa | 1 | 952 | 0 | 953 |
| AA | 0 | 0 | 256 | 256 |
| Total_HC | 24 | 953 | 256 | 1233 |

Table 84: female

| | aa | Aa | AA | Total_P3 |
|---|---|---|---|---|
| aa | 859 | 0 | 0 | 859 |
| Aa | 0 | 351 | 0 | 351 |
| AA | 0 | 1 | 60 | 61 |
| Total_HC | 859 | 352 | 60 | 1271 |

Note1. Row (Phase 3 GRCh37); Column (High Coverage GRCh38)
Note2. ‘a’: REF; ‘A’: ALT

Region: PAR2
rs number (GRCh37 phase3): .
SNP position (GRCh37) = 154954591 ; Female-Male Sex Difference in MAF (GRCh37 phase3) = -0.408 ; sdMAF P-value < 1e-300
SNP position (GRCh38) = 155724929 ; Female-Male Sex Difference in MAF (High Coverage) = -0.4088 ; sdMAF P-value < 1e-300

Table 85: REF/ALT

| | REF | ALT |
|---|---|---|
| Phase3 (GRCh37) | C | G |
| High Coverage | C | G |

Table 86: male

| | aa | Aa | AA | Total_P3 |
|---|---|---|---|---|
| aa | 22 | 1 | 0 | 23 |
| Aa | 0 | 952 | 0 | 952 |
| AA | 0 | 0 | 258 | 258 |
| Total_HC | 22 | 953 | 258 | 1233 |

Table 87: female

| | aa | Aa | AA | Total_P3 |
|---|---|---|---|---|
| aa | 855 | 1 | 0 | 856 |
| Aa | 0 | 354 | 0 | 354 |
| AA | 0 | 2 | 59 | 61 |
| Total_HC | 855 | 357 | 59 | 1271 |

Note1. Row (Phase 3 GRCh37); Column (High Coverage GRCh38)
Note2. ‘a’: REF; ‘A’: ALT

Region: PAR2
rs number (GRCh37 phase3): .
SNP position (GRCh37) = 154957867 ; Female-Male Sex Difference in MAF (GRCh37 phase3) = -0.4088 ; sdMAF P-value < 1e-300
SNP position (GRCh38) = 155728205 ; Female-Male Sex Difference in MAF (High Coverage) = -0.4088 ; sdMAF P-value < 1e-300

Table 88: REF/ALT

| | REF | ALT |
|---|---|---|
| Phase3 (GRCh37) | C | A |
| High Coverage | C | A |

Table 89: male

| | aa | Aa | AA | Total_P3 |
|---|---|---|---|---|
| aa | 22 | 0 | 0 | 22 |
| Aa | 0 | 952 | 0 | 952 |
| AA | 0 | 0 | 259 | 259 |
| Total_HC | 22 | 952 | 259 | 1233 |

Table 90: female

| | aa | Aa | AA | Total_P3 |
|---|---|---|---|---|
| aa | 857 | 1 | 0 | 858 |
| Aa | 0 | 350 | 0 | 350 |
| AA | 0 | 1 | 62 | 63 |
| Total_HC | 857 | 352 | 62 | 1271 |

Note1. Row (Phase 3 GRCh37); Column (High Coverage GRCh38)
Note2. ‘a’: REF; ‘A’: ALT

Region: PAR2
rs number (GRCh37 phase3): .
SNP position (GRCh37) = 154969038 ; Female-Male Sex Difference in MAF (GRCh37 phase3) = -0.3985 ; sdMAF P-value < 1e-300
SNP position (GRCh38) = 155739376 ; Female-Male Sex Difference in MAF (High Coverage) = -0.3973 ; sdMAF P-value < 1e-300

Table 91: REF/ALT

| | REF | ALT |
|---|---|---|
| Phase3 (GRCh37) | T | C |
| High Coverage | T | C |

Table 92: male

| | aa | Aa | AA | Total_P3 |
|---|---|---|---|---|
| aa | 21 | 2 | 0 | 23 |
| Aa | 1 | 926 | 2 | 929 |
| AA | 0 | 0 | 281 | 281 |
| Total_HC | 22 | 928 | 283 | 1233 |

Table 93: female

| | aa | Aa | AA | Total_P3 |
|---|---|---|---|---|
| aa | 812 | 4 | 0 | 816 |
| Aa | 0 | 384 | 2 | 386 |
| AA | 0 | 0 | 69 | 69 |
| Total_HC | 812 | 388 | 71 | 1271 |

Note1. Row (Phase 3 GRCh37); Column (High Coverage GRCh38)
Note2. ‘a’: REF; ‘A’: ALT

Region: PAR2
rs number (GRCh37 phase3): .
SNP position (GRCh37) = 154971425 ; Female-Male Sex Difference in MAF (GRCh37 phase3) = -0.3978 ; sdMAF P-value < 1e-300
SNP position (GRCh38) = 155741763 ; Female-Male Sex Difference in MAF (High Coverage) = -0.3974 ; sdMAF P-value < 1e-300

Table 94: REF/ALT

| | REF | ALT |
|---|---|---|
| Phase3 (GRCh37) | A | C |
| High Coverage | A | C |

Table 95: male

| | aa | Aa | AA | Total_P3 |
|---|---|---|---|---|
| aa | 37 | 0 | 0 | 37 |
| Aa | 0 | 931 | 1 | 932 |
| AA | 0 | 0 | 264 | 264 |
| Total_HC | 37 | 931 | 265 | 1233 |

Table 96: female

| | aa | Aa | AA | Total_P3 |
|---|---|---|---|---|
| aa | 841 | 2 | 0 | 843 |
| Aa | 0 | 361 | 1 | 362 |
| AA | 0 | 1 | 65 | 66 |
| Total_HC | 841 | 364 | 66 | 1271 |

Note1. Row (Phase 3 GRCh37); Column (High Coverage GRCh38)
Note2. ‘a’: REF; ‘A’: ALT

Region: PAR2
rs number (GRCh37 phase3): .
SNP position (GRCh37) = 154977261 ; Female-Male Sex Difference in MAF (GRCh37 phase3) = -0.3584 ; sdMAF P-value = 3.6584742279397e-212
SNP position (GRCh38) = 155747599 ; Female-Male Sex Difference in MAF (High Coverage) = -0.3553 ; sdMAF P-value = 2.97098408770581e-205

Table 97: REF/ALT

| | REF | ALT |
|---|---|---|
| Phase3 (GRCh37) | T | C |
| High Coverage | T | C |

Table 98: male

| | aa | Aa | AA | Total_P3 |
|---|---|---|---|---|
| aa | 28 | 0 | 0 | 28 |
| Aa | 2 | 830 | 14 | 846 |
| AA | 0 | 11 | 348 | 359 |
| Total_HC | 30 | 841 | 362 | 1233 |

Table 99: female

| | aa | Aa | AA | Total_P3 |
|---|---|---|---|---|
| aa | 675 | 9 | 0 | 684 |
| Aa | 5 | 461 | 7 | 473 |
| AA | 0 | 2 | 112 | 114 |
| Total_HC | 680 | 472 | 119 | 1271 |

Note1. Row (Phase 3 GRCh37); Column (High Coverage GRCh38)
Note2. ‘a’: REF; ‘A’: ALT
